# Neuronal signature of social novelty exploration in the VTA: implication for Autism Spectrum Disorder

**DOI:** 10.1101/280537

**Authors:** Sebastiano Bariselli, Hanna Hörnberg, Clément Prévost-Solié, Stefano Musardo, Laetitia Hatstatt-Burkle, Peter Scheiffele, Camilla Bellone

## Abstract

Novel stimuli attract our attention, promote exploratory behavior, and facilitate learning. Atypical habituation and aberrant novelty exploration have been related with the severity of Autism Spectrum Disorders (ASD) but the underlying neuronal circuits are unknown. Here, we report that dopamine (DA) neurons of the ventral tegmental area (VTA) promote the behavioral responses to novel social stimuli, support preference for social novelty, and mediate the reinforcing properties of novel social interaction. Social novelty exploration is associated with the insertion of calcium-permeable GluA2-lacking AMPA-type glutamate receptors at excitatory synapses on VTA DA neurons. These novelty-dependent synaptic adaptations only persist upon repeated exposure to social stimuli and sustain social interaction. Global or DA neuron-specific inactivation of the ASD risk gene *Neuroligin3* alters both social novelty exploration and the reinforcing properties of social stimuli. These behavioral deficits are accompanied by an aberrant expression of non-canonical GluA2-lacking AMPA-receptors at excitatory synapses on VTA DA neurons and an occlusion of novelty-induced synaptic plasticity. Altogether, these findings causally link impaired novelty exploration in an ASD mouse model to VTA DA circuit dysfunction.

## Introduction

From infancy, we encounter an array of diverse stimuli from the environment. Stimulus repetition can result in habituation whereas novel stimuli trigger elevated behavioral responses. Habituation and novelty detection allow us focusing attention on what is un-known, promote exploratory behavior, facilitate learning and are predictive of cognitive function later in life^1^. Several neuropsychiatric disorders are characterized by deficits in habituation and novelty exploration. In autism, young ASD patients show prolonged attention to depictions of objects but reduced attention to social stimuli^2^. Moreover, ASD patients exhibit aberrant habituation to social stimuli and reduced responses to social novelty^3,4^. Such alterations in novelty responses and habituation appear to be observed in a significant number of individuals with ASD, as they have been reported in clinical studies using diverse stimuli and read-outs^5–9^. However, the circuits and neuronal mechanisms underlying this specific aspect of the ASD phenotype remain largely unknown.

One system that may contribute to social novelty responses and habituation are dopamine (DA) neurons in the ventral tegmental area (VTA) and substantia nigra (SN). DA neurons increase their activity in response to novel environments^10^, stimuli of positive or negative value^11^, and natural rewards, such as food^12^. Interestingly, these neurons also respond to non-rewarding novel stimuli and their responses habituate when the stimulus becomes familiar^13,14^. This has led to the proposal that novelty by itself may be rewarding. Notably, several studies highlight decreased social reward processing in patients with ASD^15,16^ and these alterations have been hypothesized to precipitate further developmental consequences in social cognition and communication^17^.

In rodents, VTA DA neurons increase their activity in response to rewarding stimuli^18^, unfamiliar conspecifics or unfamiliar objects and this activity is necessary to promote social, but not object exploration^19^. Moreover, VTA DA neuron outputs modulate inter-peduncular nucleus (IPN) activity resulting in an enhanced exploration of familiar social stimuli^20^. Basal ganglia circuits, oxytocin, endocannabinoid, and dopamine signaling have been shown to regulate social reward behaviors^21–23^. Moreover, several mutant mouse strains, including oxytocin knock-out mice fail to respond to social novelty, a phenotype that could reflect social memory deficits^24–28^. However, the circuitry and synaptic plasticity events contributing to social novelty responses remain incompletely understood.

Glutamatergic synapses onto DA neurons undergo several forms of synaptic plasticity that may contribute to the modification of social interactions in response to experience. Specific synaptic adaptations have been described during development, after drug exposure, cue-reward learning, reciprocal social interactions and after repeated burst stimulation of DA neurons^18,29–32^. Furthermore, glutamatergic transmission is altered in several ASD animal models^33^, and we have recently shown that deficits in the postnatal development of excitatory transmission onto VTA DA neurons lead to sociability deficits^34^. Whether specific forms of synaptic plasticity in the VTA are induced by novelty exposure and whether aberrant plasticity associated with social novelty detection in the VTA is related to the maladaptive responses to social novel stimuli in ASD models is still largely unknown.

In this study, we parsed the response to social novel stimuli, social novel preference and the reinforcing properties of novel social stimuli as specific aspects of sociability controlled by DA neurons. We demonstrate that intact VTA DA neuron excitability is necessary to drive preference for social novelty but not for novel objects. Additionally, we developed a social novelty conditioned place preference protocol and show that VTA DA neuron function is required for social novelty-induced contextual reinforcement learning. Mice lacking the expression of the ASD-risk factor Neuroligin 3 (*Nlgn3*) exhibit aberrant social novelty and habituation processing. These phenotypes are recapitulated by VTA DA neuron-specific down-regulation of *Nlgn3*, thus providing a cell type-and circuit-based perspective on specific aspects of sociability dysfunctions in ASD. Finally, we discovered a form of novelty-induced synaptic plasticity at glutamatergic inputs onto VTA DA neurons that sustains social interactions and is impaired in *Nlgn3* KO and *Nlgn3* VTA DA knockdown mice.

## Results

### VTA DA neuron excitability controls social, but not object novelty exploration

To examine whether VTA DA neurons regulate novelty-induced exploration, we virally expressed the inhibitory DREADD (hM4Di)^35^ or mCherry in DA neurons of the VTA in adolescent mice (VTA::DAhM^4^Di: AAV5-hSyn-DIO-hM4Di-mCherry or VTA::DAmCherry: AAV5-hSyn-DIO-mCherry injected into DAT-Cre mice, Fig. 1a). Virus infusions led to mCherry expression in 50% of TH+ (Tyrosine hydroxylase, an enzyme necessary for DA synthesis) VTA neurons and in only very few (2%) of TH+ cells in the neighboring substantia nigra pars compacta (SNc; **Fig. S1a**), confirming preferential targeting of the VTA. Application of the hM4Di ligand Clozapine-n-oxide (CNO) decreased the neuronal excitability of VTA::DAhM^4^Di neurons compared to VTA::DAmCherry, measured *ex vivo*, as a reduction in the number of action potentials fired at increasing amplitude steps of current injections (**Fig. S1b**).

**Figure 1.**
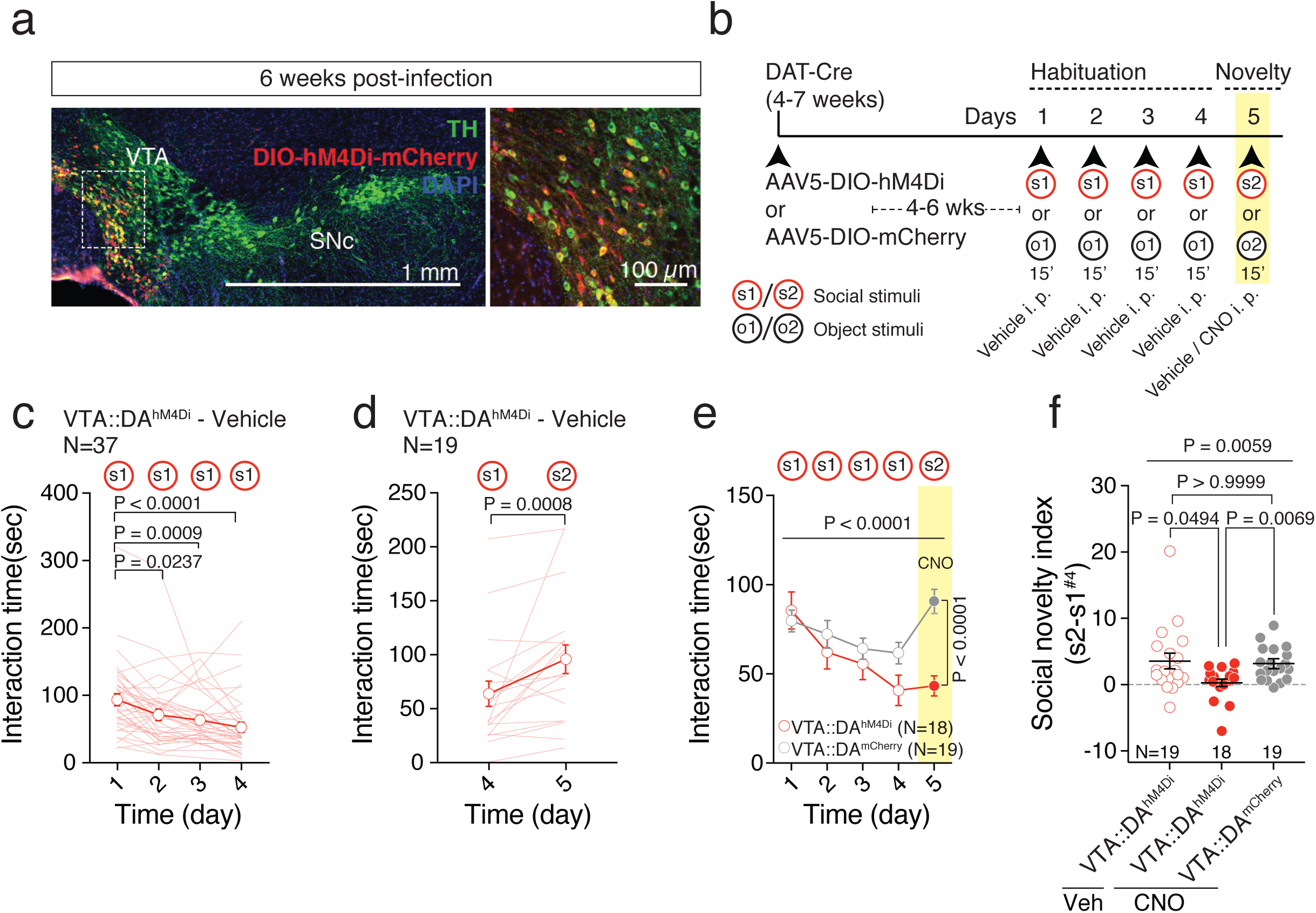
VTA DA neuron excitability controls social novelty exploration. **(a).** Representative images (low and high magnification) of immuno-staining experiments against Tyrosine Hydroxylase (TH) enzyme (in green) performed on midbrain slices of DAT-Cre mice infected with AAV5-DIO-hM4Di-mCherry (red).**(b).** Experimental time-course for the long-term habituation/novelty task. **(c)** Time course of time interaction for VTA::DAhM^4^Di mice treated with vehicle during habituation phase. Friedman test (x^2^(4) = 32.94, P < 0.0001) followed by Dunn’s test for planned multiple comparisons. **(d)** Graph reporting the time interaction at day 4 with s1 and at day 5 with s2 for VTA::DAhM^4^Di mice treated with vehicle. Wilcoxon test (W = 156). **(e)** Time interaction over days during the habituation/novelty task (s1 and s2 are social novel stimuli presented at day 1-4 and 5, respectively) for VTA::DAhM^4^Di and VTA::DAmCherry mice treated with CNO. Repeated measures (RM) two-way ANOVA (time main effect: F(4,140) = 12.38, P < 0.0001; virus main effect: F(1,35) = 3.13, P = 0.0854; time × drug interaction: F(4,140) = 9.32, P < 0.0001) followed by Bonferroni post-hoc test. **(f)** Social novelty index calculated from VTA::DAhM^4^Di treated with vehicle, VTA::DAhM^4^Di and VTA::DAmCherry both treated with CNO. Kruskal-Wallis test (K(3) = 10.26, P = 0.0059) followed by Dunn’s multiple comparisons test. N indicates number of mice. Error bars report s.e.m.

To assess the effect of the reduced excitability of DA neurons on social-novelty exploration, we conducted a social habituation/social novelty task. To enable the identification of enduring alterations in synaptic function and plasticity (see further below) we modified the classic short-term social memory paradigm^25,36^ by increasing the length of each interaction trial, performing 1 trial of 15 minutes a day over 5 days (Fig. 1b). To compare responses to social as well as non-social stimuli, we examined the behavioral responses not only for same-sex conspecifics but also objects. When repeatedly exposed to the same social (Fig. 1c) or object stimulus (**Fig. S1c**; s1 and o1, respectively), VTA::DAhM^4^Di animals injected with vehicle show reduced interaction with the stimuli over days. We refer to this process as “long-term habituation”. After 4 habituation days, the animals increased their exploratory behavior towards either a novel social stimulus (s2; Fig. 1d) or a novel object (o2, **Fig. S1d**) at day 5.

To study the role of VTA DA neurons in this behavioral trait, DA neuron excitability was decreased by intra-peritoneal (i.p.) injection of CNO in VTA::DAhM^4^Di before the exposure to the novel stimulus at day 5. We found that VTA::DAhM^4^Di animals decreased their exploratory behavior toward the new social stimulus. By contrast, VTA::DAmCherry mice treated with CNO showed unaltered novelty exploration (s2, Fig. 1e, f). Interestingly, when exposed to a novel object (o2), both VTA::DAhM^4^Di and VTA::DAmCherry animals treated with CNO exhibited novelty exploration (**Fig. S1e, f**). Thus, reducing VTA DA neuron excitability alters social novelty-induced increases in exploratory behavior but does not affect the investigation of an unfamiliar object, suggesting a differential requirement of DA neuron activity for driving exploration of social and inanimate stimuli.

### Intact VTA DA neuron excitability is necessary for preference for social novelty

When comparing stimuli of different nature, ASD patients show reduced attention to social stimuli but pay attention to non-social stimuli^37^. To assess the role of VTA DA neuron excitability in mediating both the orienting toward and the exploration of a novel social stimulus over an inanimate object or a familiar social stimulus, VTA::DAhM^4^Di and VTA::DAmCherry mice were subject to the 3-chamber test^38^ under vehicle and CNO conditions. To this end, the test was performed twice: first, the animals received either vehicle or CNO and, after 1 week of washout, the test was repeated and the pharmacological treatment was counterbalanced (Fig. 2a). To monitor potential off target effects of CNO^39^ we also included VTA::DAmCherry mice treated with CNO as controls. During the task, the animals were given a choice between an object (o1) *versus* an unfamiliar mouse (s1 or s3; social preference, SP) and subsequently a choice between a familiar (second exposure to s1 or s3) *versus* an unfamiliar social stimulus (s2 or s4; preference for social novelty, SN). Previous studies define sociability in this assay as longer time spent in the chamber with the same-sex target mouse rather than in the chamber with the object, and more time spent sniffing the same-sex mouse rather than sniffing the object^40,41^.

**Figure 2.**
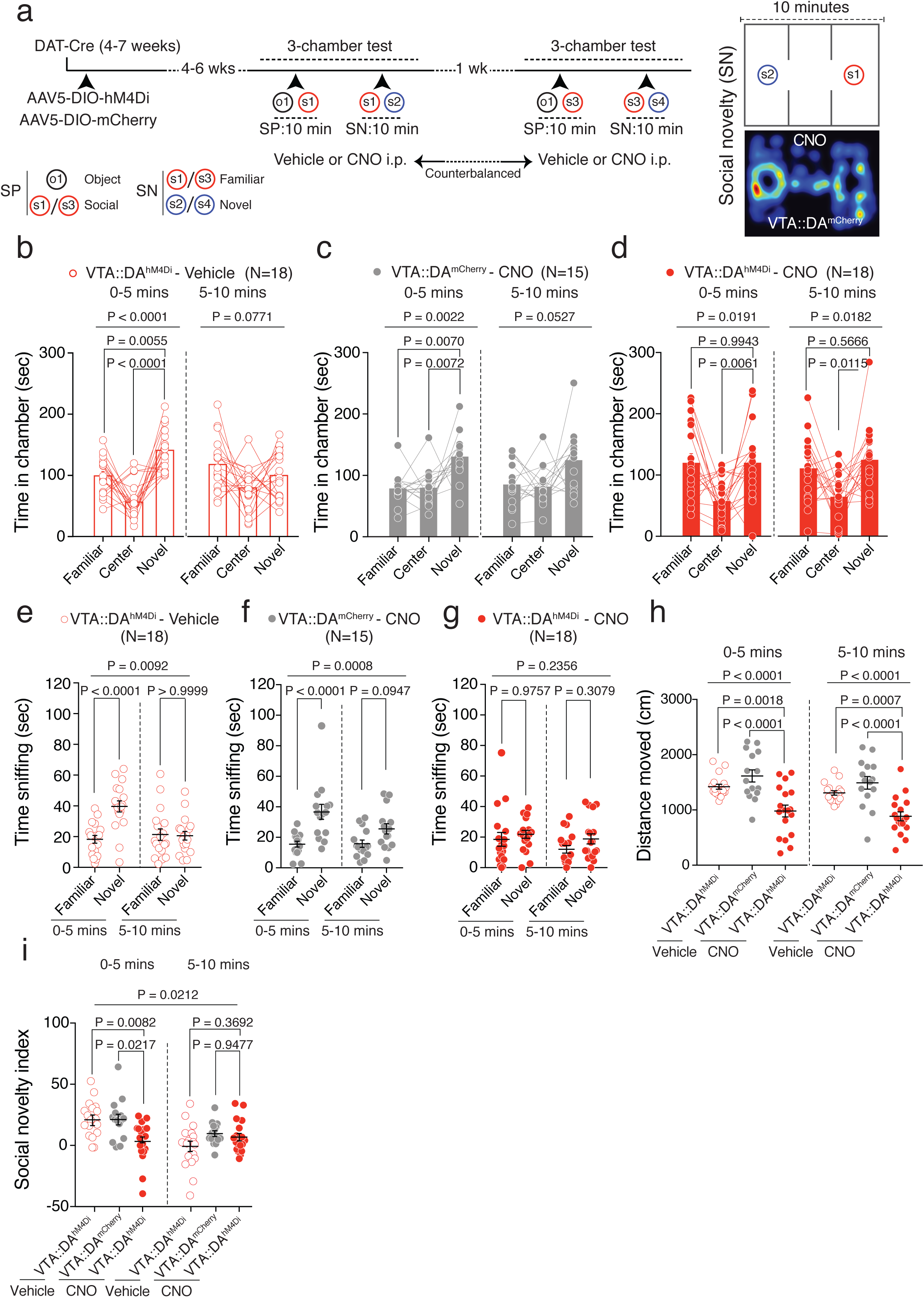
VTA DA neuron excitability controls preference for social novelty. **(a)** Left: experimental time-course for the 3-chamber test. CNO and vehicle treatments were counterbalanced. Right: schematic and occupancy plot for the 3-chamber test during social novelty phase (SN) of a CNO treated VTA::DAmCherry animal. **(b)** Time in different stimuli chamber of vehicle treated VTA::DAhM^4^Di mice. RM one-way ANOVA binned in 5 mins (chamber main effect: F(1.969, 33.48) = 21.79, P < 0.0001, first 5 mins; chamber main effect: F(1.881, 31.98) = 2.825, P = 0.0771, last 5 mins) followed by Holm-Sidak post-hoc test for planned comparisons. **(c)** Time in different stimuli chamber of CNO treated VTA::DAmCherry mice. RM one-way ANOVA binned in 5 mins (chamber main effect: F(1.645, 23.03) = 8.959, P = 0.0022, first 5 mins; chamber main effect: F(1.545, 21.63) = 3.665, P = 0.0527, last 5 mins) followed by Holm-Sidak post-hoc test for planned comparisons. **(d)** Time in different stimuli chamber of CNO treated VTA::DAhM^4^Di mice. RM one-way ANOVA binned in 5 mins (chamber main effect: F(1.494, 25.4) = 5.248, P = 0.0191, first 5 mins; chamber main effect: F(1.663, 28.27) = 5.006, P = 0.0182, last 5 mins) followed by Holm-Sidak post-hoc test for planned comparisons. **(e)** Time sniffing the novel or familiar social stimuli of vehicle treated VTA::DAhM^4^Di mice. RM two-way ANOVA (Stimulus main effect: F(1,34) = 7.634, P = 0.0092; time main effect: F(1,34) = 9.617, P = 0.0039; time × stimulus interaction: F(1,34) = 18.41, P = 0.0001) followed by Bonferroni post-hoc test. **(f)** Time sniffing the novel or familiar social stimuli of CNO treated VTA::DAmCherry mice. RM two-way ANOVA (stimulus main effect: F(1,28) = 13.96, P = 0.0008; time main effect: F(1,28) = 5.028, P = 0.0330; time × stimulus interaction: F(1,28) = 5.629, P = 0.0248) followed by Bonferroni post-hoc test. **(g)** Time sniffing the novel or familiar social stimuli of CNO treated VTA::DAhM^4^Di mice. RM two-way ANOVA (Stimulus main effect: F(1,34) = 1.458, P = 0.2356; Time main effect: F(1,34) = 4.806, P = 0.0353; time × stimulus interaction: F(1,34) = 0.6434, P = 0.4281) followed by Bonferroni post-hoc test. **(h)** Distance moved during the social novelty phase, binned in 5 mins, for VTA::DAhM^4^Di vehicle and CNO treated mice, and VTA::DAmCherry CNO treated mice. One-way ANOVA (group main effect: F(2, 48) = 12.86, P < 0.0001, first 5 mins; group main effect: F(2, 48) = 15.01, P < 0.0001, last 5 mins) followed by Bonferroni post-hoc test for planned comparisons. **(i)** Social novelty index calculated as the difference between the time sniffing the novel social stimulus minus the time sniffing the familiar social stimulus. Repeated measures two-way ANOVA (time main effect: F(2,48) = 10.54, P = 0.0021; group main effect: F(2,48) = 35.503, P = 0.0212; time × group interaction: F(2,48) = 6.23, P = 0.0039) followed by Bonferroni post-hoc test for planned comparisons. N indicates number of mice. Error bars represent s.e.m.

According to these criteria, VTA::DAhM^4^Di mice treated with vehicle (**Fig. S2a, d**), VTA::DAmCherry mice treated with CNO (**Fig. S2b, e**) as well as in VTA::DAhM^4^Di mice treated with CNO (**Fig. S2c, f**) exhibited sociability. In the social preference phase of the assay, we observed a decreased distance moved upon CNO-mediated reduction of DA neuron excitability (**Fig. S2g**). However, despite the reduced locomotion, experimental subjects still expressed social preference.

During the second phase of the 3-chamber task, preference for social novelty was defined as longer time spent in the chamber with the same-sex novel mouse rather than in the chamber with the familiar mouse and more time spent sniffing the same-sex novel mouse rather than sniffing the familiar mouse, particularly during the first 5 minutes of the test^38^. While preference for social novelty was exhibited by VTA::DAhM^4^Di mice treated with vehicle (Fig. 2b, e) and by VTA::DAmCherry mice treated with CNO (Fig. 2c, f), it was absent in VTA::DAhM^4^Di mice treated with CNO (Fig. 2d, g). As for the social preference phase, CNO treated VTA::DAhM^4^Di mice displayed a reduction in distance moved (Fig. 2h). Additionally, to compare preference for social novelty across groups, we calculated a preference score, named here “social novelty index”, as time spent sniffing the novel social stimulus *minus* time spent exploring the familiar target^42^, in the first and last 5 minutes of the assay. We found that the social novelty index was reduced by CNO injections in VTA::DAhM^4^Di mice compared to both CNO treated VTA::DAmCherry and vehicle treated VTA::DAhM^4^Di (Fig. 2i). Altogether, these findings indicate that reducing the excitability of DA neurons decreases preference for novel social stimuli when given a choice to explore either a familiar or a novel social stimulus.

### VTA DA neuron excitability mediates social novelty reinforcing properties

To investigate whether novel stimuli have reinforcing properties in mice, we designed a novelty conditioned place preference (nCPP) protocol. The protocol modifies previously used social-CPP paradigm^21,43^ as follows: mice are housed jointly with familiar mice throughout the protocol and are conditioned with novel stimuli in a test apparatus. After the Pre-TEST, we performed 4 days of repeated conditioning where wild-type (WT) mice learn to associate one compartment of the apparatus with the presence of either a co-housed social (familiar, f1), a novel social (s1) or a novel object (o1) stimulus while the other compartment is left empty (Fig. 3a, b). At day 5 (Post-TEST) the preference of mice to explore the two compartments, in the absence of social or object stimuli, was quantified and compared to Pre-TEST. While with this nCPP protocol no significant preference was developed with the familiar social stimulus (Fig. 3c and **Fig. S3a**), mice exhibited preference to explore the compartment associated with the novel social stimulus (Fig. 3d and **Fig. S3b**), and an avoidance for the novel object stimulus associated chamber (Fig. 3e and **Fig. S3c**). Interestingly, across conditioning sessions, we observed habituation to all the stimuli (Fig. 3f-h). However, when the time of interaction with the stimulus during the first and the last day of conditioning were plotted, we observed a higher interaction with novel social stimulus compared to the other stimuli at both time points (Fig. 3i). These data suggest that a novel social stimulus remains salient over days and promotes contextual associative learning.

**Figure 3.**
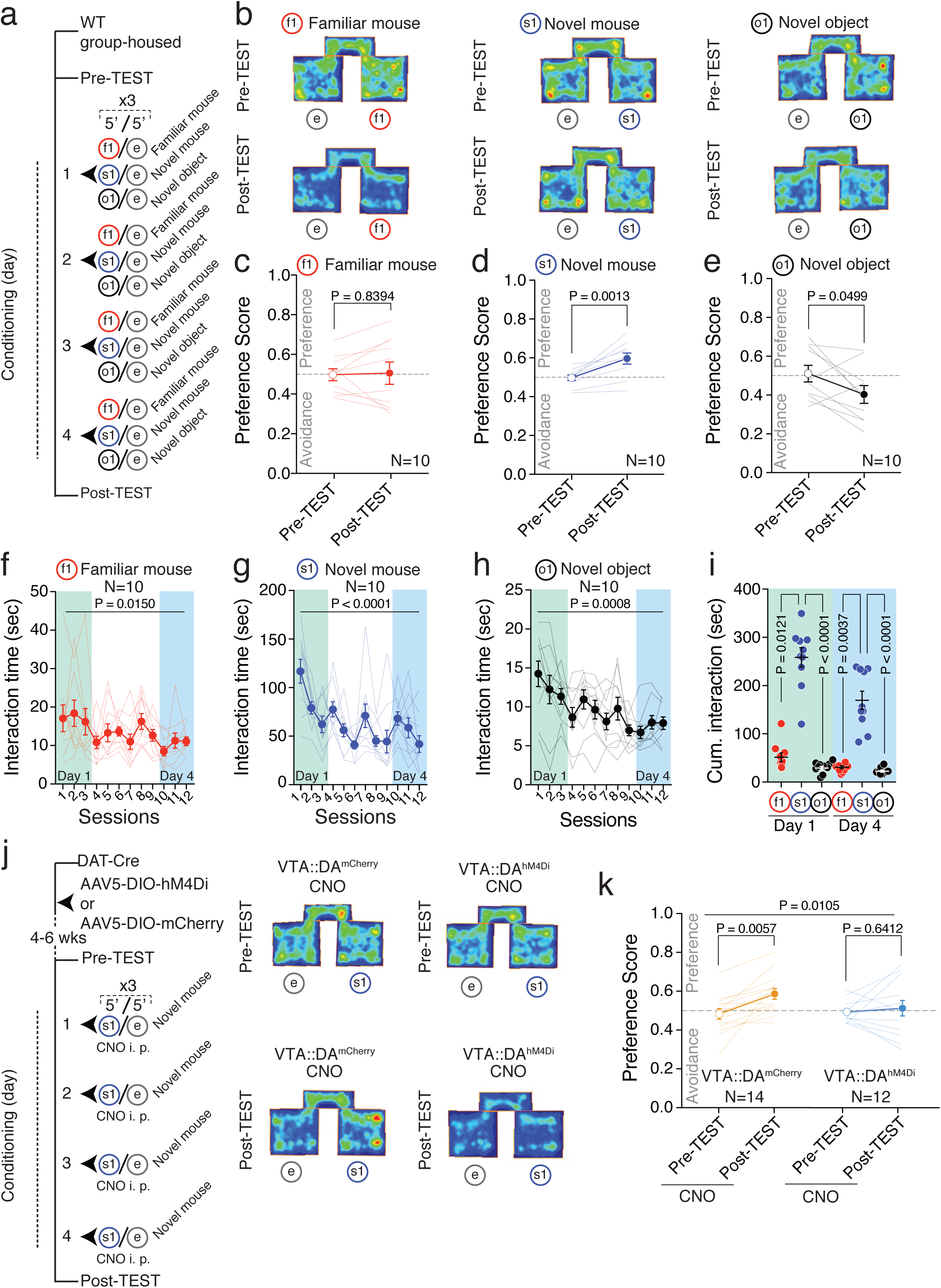
VTA DA neuron excitability mediates the reinforcing properties of social novelty. **(a)** Experimental protocol for nCPP with different stimuli. **(b)** Representative occupancy plots during Pre-and Post-TEST for mice subjected to familiar (f1), novel mouse (s1) and novel object (o1) pairing. **(c)** Scatter plot of preference score measured during the Pre-and Post-TEST for familiar mouse pairing during CPP. Paired t-test (t(9) = 0.2086; mean and s.e.m for Pre-TEST: 0.498 ± 0.0298; mean and s.e.m for Post-TEST: 0.506 ± 0.0562). **(d)** Scatter plot of preference score measured during the Pre-and Post-TEST for novel mouse pairing during CPP. Paired t-test (t(9) = 4.578; mean and s.e.m for Pre-TEST: 0.497 ± 0.0144; mean and s.e.m for Post-TEST: 0.596 ± 0.0285). **(e)** Scatter plot of preference score measured during the Pre-and Post-TEST for novel object pairing during CPP. Paired t-test (t(9) = 2.263; mean and s.e.m for Pre-TEST: 0.510 ± 0.0430; mean and s.e.m for Post-TEST: 0.403 ± 0.0455). **(f)** Time course of interaction during conditioning blocks with a familiar mouse (f1). Friedman test (P = 0.0150; x^2^(12) = 23.51). **(g)** Time course of interaction during conditioning blocks with a novel mouse (s1). Friedman test (P < 0.0001; x^2^(12) = 52.71). **(h)** Time course of interaction during conditioning blocks with a novel object (o1). Friedman test (P = 0.0008; x^2^(12) = 31.88). **(i)** Cumulative interaction with familiar mouse (f1), novel mouse (s1) and novel object (o1) during conditioning sessions at day 1 and day 4 respectively. Kruskal-Wallis test (K(6) = 46.09, P < 0.0001) followed by Dunn’s test for planned comparisons. **(j)** Left: experimental protocol for VTA::DAhM^4^Di and VTA::DAmCherry treated with CNO during CPP with novel mouse (s1) pairings. Right: representative occupancy plots during Pre-and Post-TEST for VTA::DAmCherry and VTA::DAhM^4^Di treated with CNO. **(k)** Scatter plot of preference score measured during the Pre-and Post-TEST for VTA::DAmCherry treated with CNO during conditioning sessions with a novel mouse (s1) (mean and s.e.m for Pre-TEST: 0.4808 ± 0.0267; mean and s.e.m for Post-TEST: 0.5836 ± 0.0275), and scatter plot of preference score measured during the Pre-and Post-TEST for VTA::DAhM^4^Di treated with CNO during conditioning sessions with a novel mouse (s1) (mean and s.e.m for Pre-TEST: 0.4903 ± 0.0162; mean and s.e.m for Post-TEST: 0.5091 ± 0.0428). RM two-way ANOVA (time main effect: F(1, 24) = 7.7048, P = 0.0105; virus main effect: F(1,24) = 0.8678, P = 0.3609; time × virus interaction: F(1,24) = 3.2861, P = 0.0824) followed by Bonferroni post-hoc test for planned comparisons. N indicates number of mice. Error bars represent s.e.m.

To assess the role of VTA DA neuron excitability in mediating reinforcing properties of novel social stimuli, both control VTA::DAmCherry and VTA::DAhM^4^Di received injections of CNO before each conditioning session and were treated with vehicle before the Post-TEST (Fig. 3j). Control VTA::DAmCherry but not VTA::DAhM^4^Di mice developed a preference for the compartment associated with novel social stimulus exposures (Fig. 3k and **Fig. S3d, e**). These observations suggest that the excitability of DA neurons mediates both the interaction with a novel social stimulus as well as its reinforcing properties.

### Global *Nlgn3KO* mice exhibit reduced social novelty and social reward response

Patients with ASD show aberrant responses to social novelty^4^ and are less responsive to social reward^15^. Thus, we tested whether an ASD-associated mutation in *Nlgn3*^44–46^, a gene encoding a post-synaptic adhesion molecule^47^, might result in deficits in social novelty exploration and social reward in mice as a consequence of aberrant VTA DA function. Global *Nlgn3KO* mice^48^ exhibit reduced ultrasonic vocalization and social memory in male-female interactions as well as altered motor behaviors and olfaction^26,49–51^. We examined male-male interactions in the social habituation and social novelty exploration test (Fig. 4a). *Nlgn3KO* mice exhibited overall lower interaction times, no significant habituation, and lacked the increased response to social novelty seen in wild-type littermates (Fig. 4b, c and **Fig. S4a-d**). However, *Nlgn3KO* mice showed normal habituation and novelty response to objects (Fig. 4d, e) and preference for novel objects in a novel object recognition task (Fig. 4f-h). This indicates that both novelty preference and memory for objects are unaltered. In addition to impaired social novelty response, *Nlgn3KO* mutants exhibit alterations in motor activity (Fig. 4i) and marble burying (Fig. 4j). In an olfactory discrimination test^52^, *Nlgn3KO* male mice showed normal response and habituation to a social odor (**Fig. S4e**). However, the mutant mice had a significantly decreased response when subsequently presented to a second (novel) social odor (**Fig. S4e**). To further examine social interaction in *Nlgn3KO* mice, we tested the reinforcing properties of social interaction. In a social-CPP test where the mice are conditioned with familiar mice and kept in isolation between conditioning sessions^21,43^, *Nlgn3KO* mice did not develop a preference for the social compartments whereas wild-type mice did (Fig. 4k, l, and **Fig. S4f, g**). These findings suggest that *Nlgn3KO* mice exhibit altered social interactions and defects in social reward behaviors.

**Figure 4.**
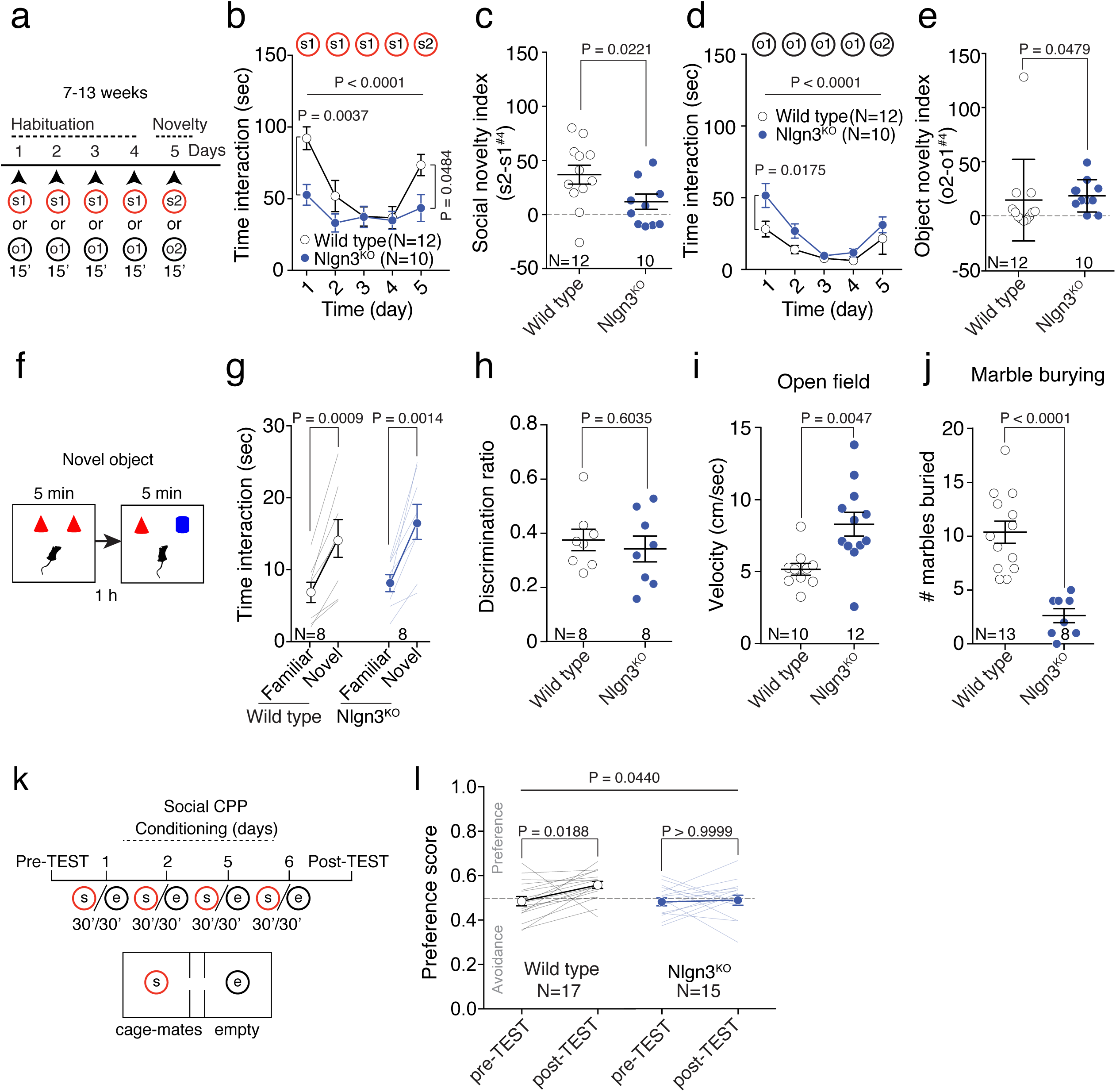
Global knockdown of *Nlgn3* alters social novelty and social reward response. (**a**) Schematic of the novelty exploration test where a mouse interacts repeatedly for 15 min with a social stimulus mouse (s1) or object (o1) for 4 days followed by a novel social stimulus mouse (s2) or novel object (o2) on the 5th day. (**b**) Mean social interaction time with the stimulus mice plotted for wild type (WT) and *Nlgn3KO* mice. RM two-way ANOVA (time main effect: F(4, 80) = 20.3, P < 0.0001; genotype main effect: F(1, 20) =3.629, P = 0.0713; time × genotype interaction: F(4, 80) = 6.071, P = 0.0003) followed by Bonferroni’s post-hoc test. (**c**) Social novelty index plotted for WT and *Nlgn3KO* mice. Unpaired t-test (t(20) = 2.481). (**d**) Mean object interaction plotted for WT and *Nlgn3KO* mice. RM two-way ANOVA (time main effect: F(4, 80) = 17.07, P < 0.0001; genotype main effect: F(1, 20) = 3.858, P = 0.0636; time genotype interaction: F(4, 80) = 1.715, P = 0.1547) followed by Bonferroni’s post-hoc test. (**e**) Dot plot of object novelty index for WT and Nlgn3KO mice. Mann-Whitney U=30. (**f**) Schematic of novel object recognition test. (**g**) Time spent investigating a novel and a familiar object during a 5-min trial 1 hour after acquisition. Paired t-test (WT: t(7) = 5.494. Mean and s.e.m familiar = 6.841 ± 1.41, mean and s.e.m novel = 14.33 ± 2.617. KO: t(7) = 5.12. Mean and s.e.m familiar = 8.115 ± 1.186, mean and s.e.m novel = 16.63 ± 2.441). (**h**) Object discrimination ratio for WT and *Nlgn3KO* mice. Unpaired t-test (t(14) = 0.5314). (**i**) Mean velocity of WT and *Nlgn3KO* mice during a 7 min open field test. Unpaired t-test (t(20) = 3.178). (**j**) Number of marbles (out of 20) buried during a 30 min marble burying task plotted for WT and *Nlgn3KO* mice. Unpaired t-test (t(19) = 5.505). (**k**) Schematics of the social conditioned place preference (CPP) test. (**l)** Scatter plot of preference score measured during the Pre-and Post-TEST for WT (mean and s.e.m for Pre-TEST: 0.4846 ± 0.0209; mean and s.e.m for Post-TEST: 0.5578 ± 0.0158), and *Nlgn3KO* mice (mean and s.e.m for Pre-TEST: 0.4809 ± 0.0178; mean and s.e.m for Post-TEST: 0.4886 ± 0.0225). RM two-way ANOVA (time main effect: F(1, 30) = 4.422, P = 0.0440; genotype main effect: F(1,30) = 3.492, P = 0.0715; time × genotype interaction: F(1,30) = 2.885, P = 0.0998) followed by Bonferroni post-hoc test for planned comparisons. N numbers indicate mice. All error bars are s.e.m.

### *Nlgn3* in VTA DA neurons is required for social novelty and social interaction

The diverse alterations in social but also non-social behaviors in *Nlgn3KO* mice, indicate that multiple different systems might contribute to their phenotype. To test whether any alterations are due to *Nlgn3* functions in VTA DA neurons we generated microRNA-based knock-down vectors for conditional suppression of *Nlgn3* expression (**Fig. S5a, b**). Cre-dependent AAV-based vectors were injected into the developing VTA of DAT-Cre mice at postnatal days 5-6 and mice were analyzed using a battery of behavioral tests (AAV2-DIO-*miRNlgn*^*3*^ in DAT-Cre mice: VTA::DANL^3^KD, Fig. 5a, b, and see **Fig. S5c** for off-target areas affected and **Fig S5d, e** for further controls). Notably, VTA::DANL^3^KD mice exhibited a similar impairment in social-CPP as the global *Nlgn3KO* mice (Fig. 5c, d, and **Fig. S5f, g**) indicating that *Nlgn3* downregulation in VTA DA neurons is sufficient to mimic this aspect of the global *Nlgn3KO* phenotype. Furthermore, in the social habituation/novelty exposure test VTA::DANL^3^KD mice showed an overall reduction in social exploration and a blunted response to novelty (Fig. 5e-g, and **Fig. S5h-k**). At the same time, VTA::DANL^3^KD mice showed preference to novel objects in the novel object task (Fig. 5h-j). Thus, there is a specific requirement for *Nlgn3* in VTA DA neurons for appropriate social novelty responses and for the reinforcing properties of social interaction. By contrast, motor activity, marble burying, and social olfaction that are altered in global *Nlgn3KO* mice were not modified in the VTA::DANL^3^KD mutants (Fig. 5k, l, **Fig. S5l**). Interestingly, we observed that knock-down of *Nlgn3* in VTA-DA neurons of adult mice produced a similar but less pronounced social interaction phenotype as in developing animals, with reduced habituation and reduced social novelty response (**Fig. S6**). Thus, *Nlgn3* is required for normal function and/or plasticity in VTA DA cells, even in fully developed circuits.

**Figure 5.**
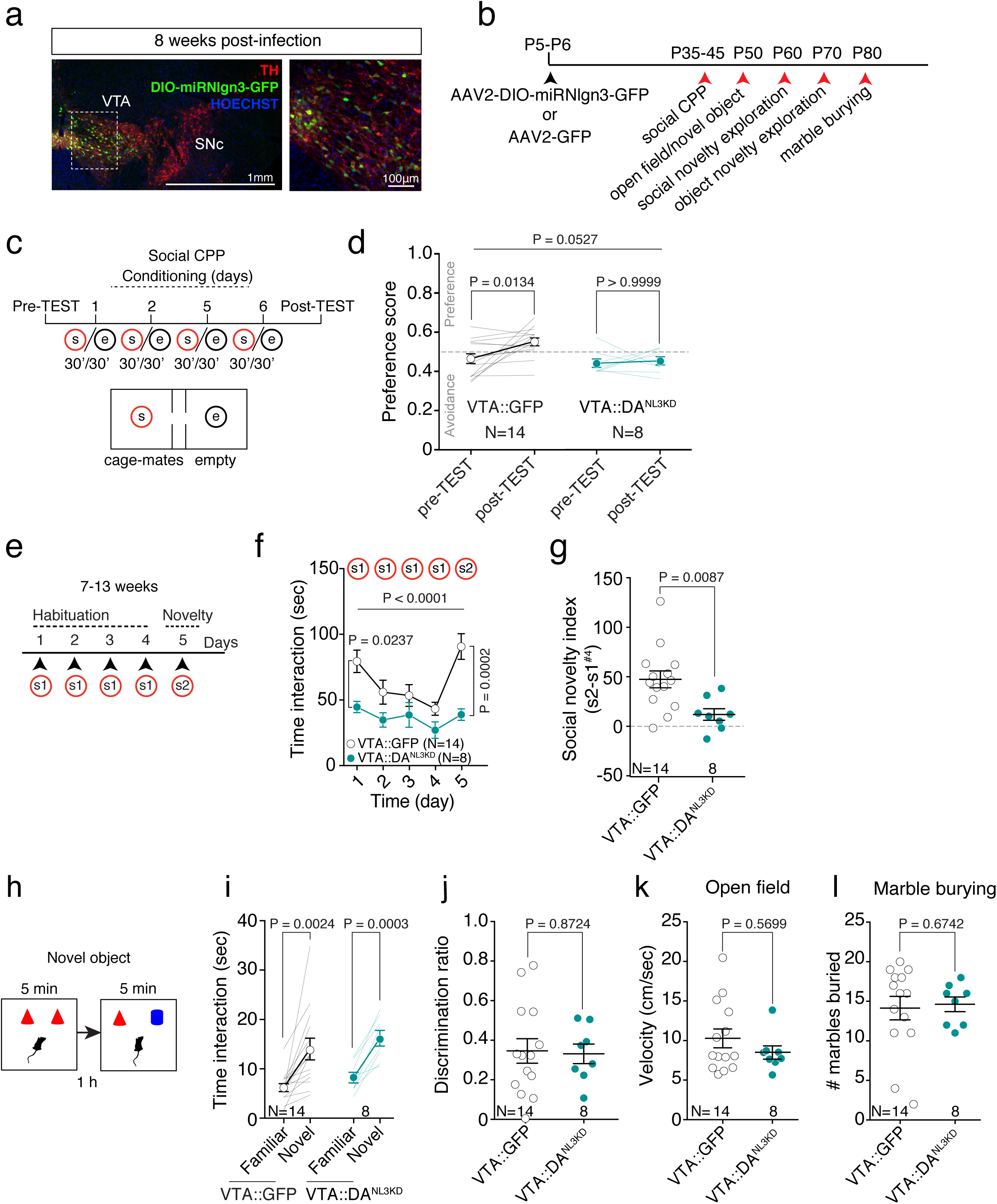
*Nlgn3* in VTA DA neurons is required for social novelty responses and social interaction. (**a)** Left: representative image of coronal slice of VTA and SNc from an AAV2 DIO-miRNlgn^3^-GFP infected DAT-Cre mouse. Right: higher magnification of VTA. (**b**) Experimental schematic of behavioral test order in VTA-injected mice. (**c**) Experimental schematic of the social-CPP test. (**d**) Scatter plot of preference score measured during the Pre-and Post-TEST for VTA::GFP (mean and s.e.m for pre-TEST = 0.4642 ± 0.0247. Mean and s.e.m post-TEST = 0.5526 ± 0.0200), and VTA::DANL^3^KD mice (mean and s.e.m for Pre-TEST: 0.4434 ± 0.0218; mean and s.e.m for Post-TEST: 0.4548 ± 0.0214). RM two-way ANOVA (time main effect: F(1, ^20^) = 4.24, P = 0.0527; virus main effect: F(1,20) = 6.103, P = 0.0226; time × viru interaction: F(1,20) = 2.527, P = 0.1276) followed by Bonferroni post-hoc test for planned comparisons. (**e**) Experimental schematic of the social novelty exploration test. (**f**) Mean social interaction plotted for VTA::GFP and VTA::DANL^3^KD mice. RM two-way ANOVA (time main effect: F(4, 80) = 8.058, P < 0.0001, virus main effect: F(1, 20) = 9.164, P = 0.0067; time × virus interaction: F(4, 80) = 3.179, P = 0.0178) followed by Bonferroni’s post-hoc test. (**g**) Social novelty index for VTA::GFP and VTA::DANL^3^KD mice. Unpaired t-test (t(20) = 2.908). (**h**) Experimental schematic of novel object recognition test. (**i**) Time spent investigating a novel and a familiar object. Paired t-test (VTA::GFP: t(13) = 3.763. Mean and s.e.m familiar = 6.199 ± 0.805, mean and s.e.m novel = 14.03 ± 2.188. VTA::DANL^3^KD: t(7) = 6.518. Mean familiar = 8.226, s.e.m ± 1.069, mean novel = 16.2, s.e.m ± 1.582). (**j**) Discrimination ratio for object discrimination plotted for VTA::GFP and VTA::DANL^3^KD. Unpaired t-test (t(20) = 0.1627). (**k**) Mean velocity of VTA::GFP and VTA::DANL^3^KD mice during a 7 min open field test. Mann-Whitney U=47. (**l**) Number of marbles buried plotted for VTA::GFP and VTA::DANL^3^KD. Mann-Whitney U=49.5. N numbers indicate mice. All error bars are s.e.m.

### A synaptic signature of saliency detection in VTA DA neurons

Several experiences strengthen synaptic transmission and drive the insertion of GluA2-lacking AMPARs at excitatory inputs onto DA neurons. This form of plasticity can be assessed by calculating a rectification index (RI) and has been observed both in the VTA^30,31^ and in dorsal raphe^53^. We used the long-term habituation/novelty task to test whether novelty exploration induced specific forms of long-lasting synaptic plasticity at excitatory inputs onto DA neurons in the VTA. In WT mice, the RI increased at synapses 24 hours after exploration of either a novel mouse or a novel object when compared to RI calculated from home caged mice (Fig. 6a). By contrast, the RI was unchanged after the exposure to a new context and AMPA/NMDA ratios were unchanged for any of the above conditions (Fig. 6b). When AMPAR EPSCs were recorded after repeated exposure (over 4 days) to object stimuli, the RI was normalized to control condition (Fig. 6c). A subsequent exposure to a new object (o2) promoted the increase in RI (**Fig. S7a**). By contrast, GluA2-lacking AMPARs were detected in mice repeatedly exposed to a social stimulus (s1) over a four-day period and were still present at these synapses after ten days of repeated exposure (Fig. 6c). Remarkably, the AMPA/NMDA ratio was significantly elevated after 4 days of social (s1) repeated exposure relative to baseline but was normalized after 10 days of repeated exposure (Fig. 6d), while the Paired-Pulse Ratio (PPR) remained unchanged throughout (**Fig. S7b**). Taken together, these data indicate that repeated exposure to a novel social stimulus, but not an object stimulus, transiently increases synaptic strength (AMPA/NMDA) and produces a stable insertion of GluA2-lacking AMPARs at VTA DA neuron excitatory inputs.

**Figure 6.**
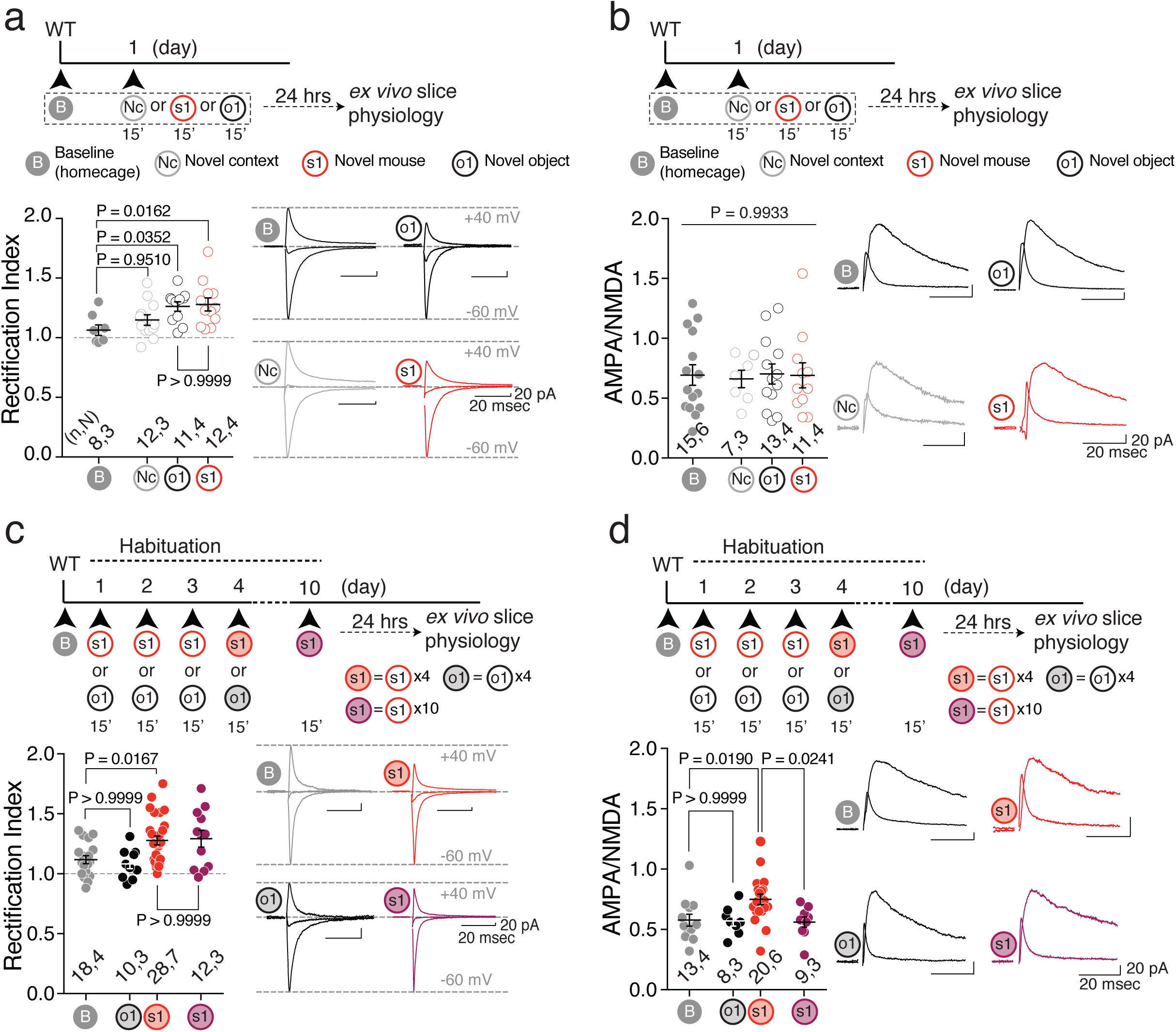
Novelty exploration-induced synaptic plasticity. **(a)** Top: experimental paradigm. Bottom: scatter plot of rectification index and AMPAR-EPSCs example traces (−60, 0 and 40 mV) recorded from VTA DA neurons at baseline (B, homecage), or 24 hours after 15 minutes of novel context (Nc), novel mouse (s1) or novel object (o1) exposure. One-way ANOVA (F(3, 39) = 4.153, P = 0.0120) followed by Bonferroni post-hoc test for planned comparisons. **(b)** Top: experimental paradigm. Bottom: scatter plot and example traces of AMPA/NMDA ratio recorded from VTA DA neurons at baseline (B, homecage), or 24 hours after 15 minutes of novel context (Nc), novel mouse (s1) or novel object (o1) exposure. One-way ANOVA (F(3, 42) = 0.0287, P = 0.9933). **(c)** Top: experimental paradigm. Bottom: scatter plot and example traces of Rectification Index recorded from VTA DA neurons at baseline (B), 24 hours after 4 repeated exposures to either a novel mouse (s1) or a novel object (o1) and 10 repeated exposures to a novel mouse (s1, bold purple). One-way ANOVA (F(3, 64) = 5.149, P = 0.0030) followed by Bonferroni post-hoc test for planned multiple comparisons. **(d)** Top: experimental paradigm. Bottom: scatter plot and example traces of AMPA/NMDA ratio recorded from VTA DA neurons at baseline (B), 24 hours after 4 repeated exposures to either a novel mouse (s1) or a novel object (o1) and 10 repeated exposures to a novel mouse (s1, bold purple). One-way ANOVA (F(3,46) = 4.4939, P = 0.0076) followed by Bonferroni post-hoc test for planned multiple comparisons. n,N indicates number of cells and mice respectively. Error bars report s.e.m.

To understand the functional role of non-canonical AMPARs inserted during repeated social novelty exposure, we infused the GluA2-lacking AMPAR blocker NASPM into the VTA starting from the second day of interaction with either social or object stimuli (Fig. 7a,b). NASPM infused mice reduced the interaction with a social stimulus upon repeated exposure (Fig. 7c); by contrast, the infusions did not alter long-term habituation to an object (Fig. 7d), social interaction in the home cage between two familiar mice or distance moved in an open field (**Fig. S7c-e**). To further understand the impact of GluA2-lacking AMPARs at VTA DA neuron inputs on social long-term habituation, we promoted the insertion of GluA2-lacking AMPARs *via* blue-light illumination of ChR2 or eYFP expressing VTA DA neurons^32^ of DAT-Cre mice (VTA::DAChR^2^: AAV5-Ef1*α*-DIO-ChR2(H134R)-eYFP, VTA::DAeYFP: AAV5-Ef1*α*-DIO-eYFP). DA neuron stimulation consisted in ChR2-mediated bursts of action potentials^34^ delivered for 15 minutes twenty-four hours before each social exposure (Fig. 7e,f). This non-contingent burst activation increased RI in photocurrent positive neurons (I +; Fig. 7g) and blocked long-term habituation to social stimuli (Fig. 7h). Altogether, these data indicate that GluA2-lacking AMPARs might represent a synaptic signature of social stimulus saliency and, once inserted, their activity counteracts habituation.

**Figure 7.**
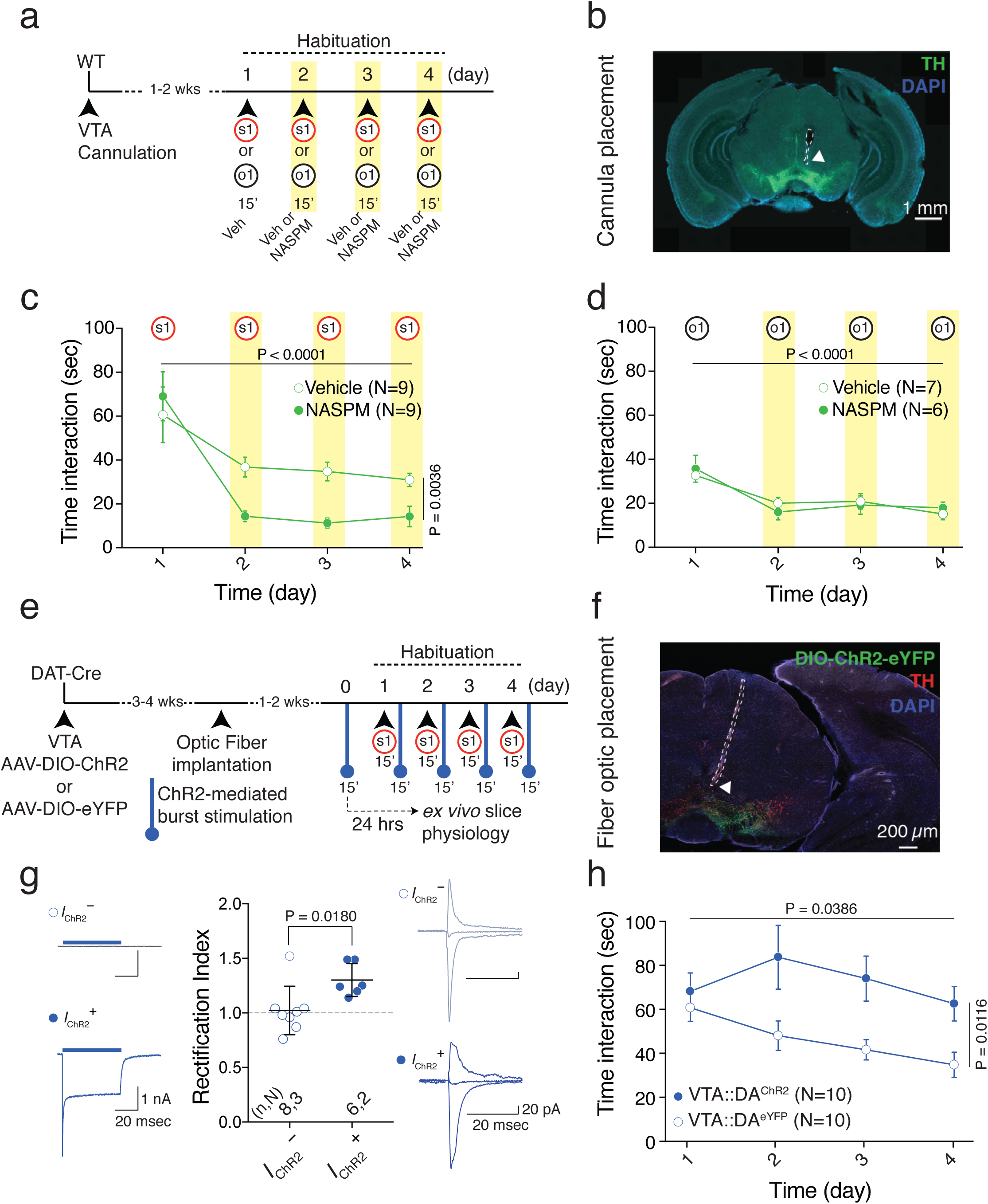
GluA2-lacking AMPAR function controls habituation to social stimuli. **(a)** Schema of the experimental paradigm. **(b)** Representative image of cannula placement for NASPM or vehicle infusion (green: TH; blue: DAPI; white arrow indicates cannula tip). **(c)** Time course of time interaction with a novel mouse (s1) for vehicle or NASPM infused mice at day 2, day 3 and day 4. RM two-way ANOVA (time main effect: F(3, 24) = 17.57, P < 0.0001; drug main effect: F(1, 8) = 16.48, P = 0.0036; time × drug interaction: F(3, 24) = 3.141, P = 0.0439). **(d)** Time course of time interaction with a novel object (o1) over 4 days for Vehicle and NASPM groups. RM two-way ANOVA (time main effect: F(3, 33) = 24.71, P < 0.0001; drug main effect: F(1,11) = 0.00005, P = 0.9942; time × drug interaction: F(3, 33) = 1.109, P = 0.3595). **(e)** Experimental paradigm for non-contingent optogenetic stimulation. **(f)** Representative image of fiber optic placement for DIO-ChR2 expressing mice (red: TH, green: AAV-DIO-ChR2-eYFP, blue: DAPI; white arrow indicates fiber optic tip). **(g)** Left: example traces of a photocurrent negative (I -) and a photocurrent positive (I +) VTA DA neuron. Middle: scatter plot of RI recorded from photocurrent negative (I -) and photocurrent positive (I +) VTA DA neurons and AMPAR-EPSCs example traces (−60, 0 and 40 mV) recorded from VTA DA neurons. Mann-Whitney test (U = 6). **(h)** Time course over 4 days of time interaction with a novel mouse (s1) for VTA::DAChR^2^ and VTA::DAeYFP mice with non-contingent optical stimulation. RM two-way ANOVA (time main effect: F(3, 18) = 2.9966, P = 0.0386; virus main effect: F(1, 18) = 7.9034, P = 0.0116; time × virus interaction: F(3, 18) = 1.9532, P = 0.1320). N indicates number of mice. Error bars represent s.e.m.

### Nlgn3 loss-of-function impairs novelty-induced plasticity

*Nlgn3* has been implicated in the regulation of AMPARs at glutamatergic synapses^49,54^. We therefore hypothesized that defects in DA neuron synaptic function could represent the mechanism underlying the aberrant habituation and response to novel social stimuli in VTA::DANL^3^KD mice. We explored glutamate receptor function in VTA DA neurons of global *Nlgn3KO* and conditional VTA::DANL^3^KD mice. Notably, we observed increased RI of AMPAR-mediated currents indicating the aberrant presence of GluA2-lacking AMPARs at excitatory inputs onto VTA DA neurons in both *Nlgn3* loss-of-function models (Fig. 8a). Given the abnormal elevation of GluA2-lacking AMPARs in naïve VTA::DANL^3^KD mice, we hypothesized that in these mice social novelty-induced plasticity might be occluded. Indeed, GluA2-lacking AMPARs in VTA DA neurons were not further increased 24 hours after novelty exposure in VTA::DANL^3^KD mice (Fig. 8b). Thus, aberrant plasticity of GluA2-lacking AMPARs in VTA DA neurons is associated with an impaired response to a social novel stimulus.

**Figure 8.**
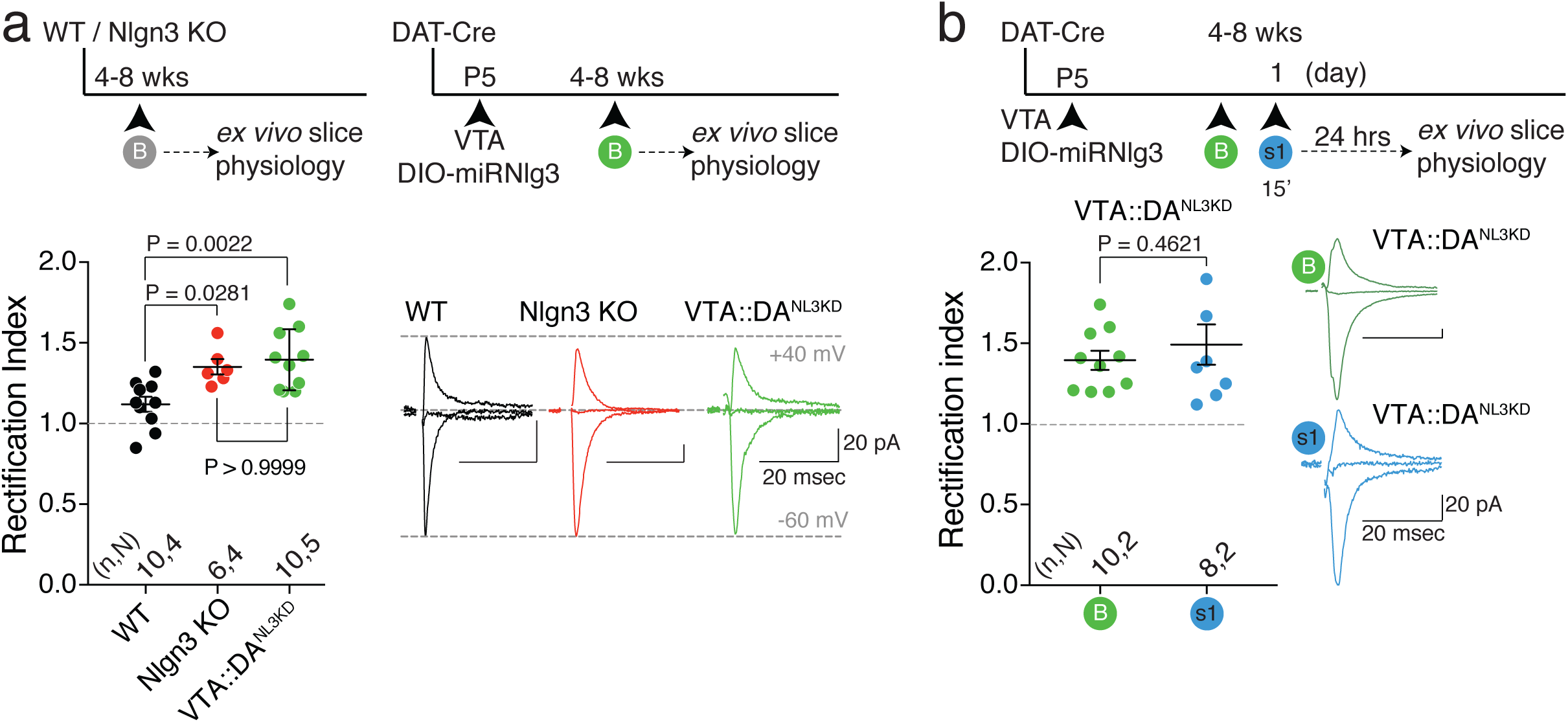
Aberrant increase of GluA2-lacking AMPARs in *Nlgn3*-deficient VTA DA neurons. **(a)** Top: experimental paradigm. Bottom: scatter plot of rectification index and example traces of AMPAR-EPSCs (−60, 0 and 40 mV) measured from adolescent WT, Nlgn3KO and VTA::DANL^3^KD. One-way ANOVA (F(2, 23) = 8.363, P = 0.0019) followed by Bonferroni post-hoc test. **(b)** Top: experimental paradigm. Bottom: scatter plot of rectification index and example traces of AMPAR-EPSCs (−60, 0 and 40 mV) measured from VTA::DANLKD mice at baseline (B) or 24 hours after 15 minutes exposure to a novel social stimulus (s1). Unpaired t-test (t(16) = 0.7536). n, N indicates number of cells and mice respectively. Error bars represent s.e.m.

## Discussion

In this study, we established that VTA DA neurons drive social novelty exploration and preference, two aspects of social behavior. Novel stimuli, independent of their nature, leave a plasticity trace at glutamatergic synapses in the VTA, which persists upon repeated exposure to social stimuli and supports sustained social interactions. We used deletion of *Nlgn3*^44–46,48^ to test whether an ASD-relevant genetic lesion might impair social novelty exploration. Global *Nlgn3* knock-out resulted in an impairment of social novelty and social reward responses. Notably, global loss of *Nlgn3* is also accompanied by a broad spectrum of additional phenotypes, including changes in olfaction and in motor-related behaviors^26,49–51^. Thus, the origin of social behavior alterations in these mice was unclear. Further, previous studies explored phenotypes in mice carrying a point mutation in *Nlgn3* that reduces (but does not abolish) *Nlgn3* expression and has been observed in 2 patients from one family ^55^. For this model, it was concluded that behavioral phenotypes are significantly dependent on the genetic context with significant phenotypes reported for some genetic backgrounds but not others ^56–60^. Here, we demonstrate that selective inactivation of *Nlgn3* in VTA DA neurons disrupts novelty-induced plasticity at glutamatergic synapses in the VTA, social novelty responses, and the reinforcing properties of social interactions while having no detectable effect on motor behaviors or olfaction. Thus, our work not only uncovers a major role for VTA DA neurons in social novelty responses but also identifies a circuit-based mechanism underlying one central aspect of a social interaction phenotype in a genetic model of ASD.

Rewarding experiences and novel stimuli are learning signals that activate VTA neurons and increase the functional connectivity between the VTA and the Nucleus Accumbens (NAc)^13,61,62^. Remarkably, the activity of DA neurons is linked to the nature of the stimulus and its saliency^63^. Social interactions are salient experiences and the novelty associated with social stimuli might promote incentive salience and contextual learning^64^. Several recent findings indicates that VTA DA neurons, with their complex input and output structures, are part of a “socially engaged reward circuit” ^65^.

Previous work demonstrated that optogenetic activation of VTA DA neurons enhances the interaction with social but not object stimuli and that stimulation of VTA DA terminals to the interpeduncular nucleus controls the expression of preference for social novelty^20^. Furthermore, medial preoptic area^65^ and paraventricular nucleus^23^ have been identified as important inputs to VTA in promoting pro-social behaviors. These studies provided instrumental information to understand the circuits within the reward system recruited during social interaction. In the present study, we identified the synaptic adaptations that occur in VTA DA neurons during social interaction and advanced our insights into the interplay between novelty and saliency processing, social reward and social interaction. Interestingly, while chemogenetic-mediated excitability reduction or conditional suppression of *Nlgn3* in VTA DA neurons affected social novelty responses, they both failed to modify responses to novel objects. The experience-dependent impact of our manipulations might result from the higher intrinsic saliency of social stimuli compared to inanimate objects. Consistent with this hypothesis, we observed that while both a novel object and a novel social stimulus triggered insertion of GluA2-lacking AMPARs at glutamatergic synapses onto VTA DA neurons. However, only the repeated exposure to novel social stimuli resulted in a transient increase of synaptic strength and the maintenance of GluA2-lacking AMPARs. Therefore, we hypothesize that while the insertion of non-canonical AMPARs at VTA DA neurons reflects the novelty associated to the stimulus, their persistence signals the higher saliency of the social over the object stimulus.

The insertion and the expression of non-canonical AMPARs is also associated with non-social, highly salient experiences. For example, cocaine and other addictive drugs promote the insertion of GluA2-lacking AMPARs at excitatory inputs onto VTA DA neurons^30,32^ and the persistence of cocaine-induced synaptic changes in the VTA affects synaptic plasticity in the downstream regions involved in cue-induced cocaine-seeking behavior^66^. However, a causal relationship between behavioral responses to salient stimuli and GluA2-lacking AMPAR expression in VTA DA neurons has not been reported. While our optogenetic and pharmacological evidence indicates that non-canonical AMPARs at VTA DA neurons contribute to behavioral responses to social stimuli, they could also represent a functionally-relevant synaptic signature responsible for the behavioral responses associated with other salient stimuli. Although cocaine administration modulates the acquisition of social reward CPP in mice and rats^67–70^, further investigations are needed to test whether persistence of GluA2-lacking AMPAR in VTA DA neurons is both a behaviorally-relevant and a shared synaptic signature of diverse salient experiences.

It has previously been reported that, in mice, social interaction with a stranger con-specific, triggers long-term potentiation in VTA DA neurons, as determined by an increase in AMPAR/NMDAR ratio^29^, a proxy of synaptic strength. Here, together with a persistent change of non-canonical AMPAR content, we uncovered upon repeated exposure to social, but not object stimuli a transient increase in AMPA/NMDA at VTA DA neuron excitatory inputs. Thus, while the insertion of GluA2-lacking AMPARs takes place in response to novelty, both the persistence of GluA2-lacking AMPARs and the transient increase in AMPA/NMDA appear to be associated to the saliency of the experience. Consistently, changes in AMPA/NMDA changes occur in response to both rewarding and aversive experiences^71^ and control associative learning^18,72^. However, whether the transient changes in synaptic strength due to repeated exposures to novel social stimuli are permissive for contextual learning warrants further investigation.

The specific synaptic signatures observed in response to social novelty responses might occur in dedicated circuits, within the DA system, responsive for processing highly salient social stimuli in a temporally-defined manner. Electrophysiological recordings from the VTA have pointed to the fact that DA neurons constitute a heterogeneous population in terms of intrinsic properties, projection specificity and neurotransmitter/neuromodulator release^73–75^. This diversity is thought to subserve the processing of reinforcing and aversive experiences in an input-output specific manner^76–78^. VTA DA neurons projecting to the nucleus accumbens (NAc), but not prefrontal cortex (PFC), control social interaction^19^ while IPN-projecting DA neurons are involved in preference for social novelty expression^20^. Therefore, the synaptic adaptations reported in response to repeated exposure to social novelty exposure might occur in DA neurons projecting to either the NAc, the IPN or both. At the same time, given the intrinsic diversity of sensory and emotional information provided by social *vs* inanimate stimuli, it is also conceivable that synaptic plasticity occurs at specific inputs to defined subclasses of VTA DA neurons. Additional investigations of synaptic properties of defined inputs to projection-specific DA neuron subclasses is needed to further understand the circuits and the synaptic mechanisms underlying both novelty and saliency processing associated with social and inanimate stimuli.

Altered social interactions and communication are defining aspects of the autism phenotype. However, such alterations may arise from a plethora of neuronal processing defects, ranging from alterations in perception, sensory processing, multisensory integration, or positive and negative valence assigned to social stimuli ^17,79,80^. In this work, we specifically explored neuronal circuitry relevant for social novelty responses. We chose this domain, as studies in children with ASD demonstrated altered habituation and responses to novel stimuli^81^, ^8^. Notably, in toddlers, a slowed habituation to faces but normal habituation to repeatedly viewed objects has been reported to coincide with more severe ASD symptoms^3^. Several rodent models of ASD exhibit altered social novelty responses^25^, ^28^, ^27^, ^82–84^ and such alterations have been suggested to reflect changes in social memory or discrimination. However, brain areas and circuit elements contributing to these changes in habituation and social novelty responses in mice and humans are largely unknown. Our rodent work not only highlights a contribution of VTA DA neurons to this process but also takes steps toward identification of the synaptic basis of social novelty responses and habituation. Considering the complexity of ASD behavioral dysfunctions, we propose that fractionating the autism phenotype according to specific behavioral domains based on neuronal circuit elements will provide a productive stratification criterion for patient populations. Thus, we speculate that in a sub-population of individuals with ASD alterations VTA DA function might contribute to the social interaction phenotype whereas in other sub-groups of patients alterations in social interaction may arise for different reasons. A prediction from this hypothesis is that stratification of patient populations based on an assessment of novelty responses, habituation, and social reward may help to identify sub-groups of patients that would particularly benefit from interventions targeting function and plasticity of the VTA-DA circuit elements.

## Acknowledgements

C.B is supported by the Swiss National Science Foundation, Pierre Mercier Foundation, and NCCR Synapsy. P.S is supported by the Swiss National Science Foundation, NCCR Synapsy, and EU-AIMS and the Canton Basel-Stadt. H.H is supported by the Human Frontier Science Program. We thank Carmen Sandi and Marie Schaer for the critical reading of the manuscript. We thank Stamatina Tzanoulinou for insights on behavioral experiments and Lorena Jourdain and Caroline Bormann for technical support.

## Author Contributions

*Ex vivo* electrophysiology experiments were performed by S.B., C.B and S.M. Behavioral experiments and analysis reported in Fig. 1, 2, 6, Sup. Fig. 1 and Sup. Fig. 2 were performed by C.P.S. Behavioral experiments and analysis of the Fig. 3 and Sup. Fig. 3 were conducted by S.B. Behavioral experiments and analysis of the Fig. 4, 5 and Sup. Fig. 4, 5 and 6 were performed by H.H. and L.H.B. S.B. and C.P.S. performed the statistical analyses for the *ex vivo* electrophysiology and the behavioral experiments in Fig. 1, 2, 3, 6 and Sup. Fig. 1, 2, 3. H.H. performed the statistical analyses for the behavioral experiments in Fig. 4, 5 and Sup. Fig. 4, 5 and 6. Generation and validation of viral knock-down vectors was performed and analyzed by H.H. Immunohistochemistry in VTA::DAhM^4^Di and cell counting were performed by C.P.S. and S.B. Immunohistochemistry and cell counting in VTA::DANL^3^KD mice were performed by H.H. The study was designed and the manuscript written by C.B., P.S., H.H. and S.B., with assistance from C.P.S.

## Material and Methods

### Animals

The study was conducted with wild-type (WT) and transgenic mice in C57BL/6J background. WT mice were obtained from Charles River. For dopamine neuron-specific manipulations DAT-iresCre (*Slc6a3tm^1^.^1^(cre)Bkmn*)^85^ and DAT-Cre BAC transgenic mice^86^ were employed. *Nlgn3KO* mice were previously described^48^. Male and female mice were housed in groups (weaning at P21 – P23) under a 12h light – dark cycle (7:00 a.m. – 7:00 p.m.). All physiology and behavior experiments were performed during the light cycle. For *Nlgn3KO* and WT mice, multiple behavioral tests were performed with the same group of animals, with a minimum of 3 days in-between tests. VTA::DANL^3^KD and VTA::GFP participated in one behavioral test prior to the start of social CPP. Embryos for cortical cultures were obtained from NMRI mice (Janvier). All the procedures performed at UNIGE and Biozentrum complied with the Swiss National Institutional Guidelines on Animal Experimentation and were approved by the respective Swiss Cantonal Veterinary Office Committees for Animal Experimentation.

### Surgery

Injections of rAAV5-hSyn-DIO-hM4D(Gi)-mCherry and rAAV5-hSyn-DIO-mCherry were performed in DAT-Cre mice at 4 – 7 weeks. Mice were anesthetized with a mixture of oxygen (1L/min) and isoflurane 3% (Baxter AG, Vienna, Austria) and placed in a stereotactic frame (Angle One; Leica, Germany). The skin was shaved, locally anesthetized with 40 – 50 µL lidocaine 0.5% and disinfected. Bilateral craniotomy (1 mm in diameter) was then performed over the VTA at following stereotactic coordinates: ML ±0.5 mm, AP –3.2 mm, DV –4.20±0.05 mm from Bregma. The virus was injected via a glass micropipette (Drummond Scientific Company, Broomall, PA) into the VTA at the rate of 100 nl/min for a total volume of 200 nL in each side. The virus was incubated for 3 – 8 weeks prior to perform the behavioral tasks or electrophysiological recordings.

Injections of purified AAV2-DIO-miRNlgn3-GFP, AAV2-Synaptophysin-GFP and AAV2-DIO-miR-GFP were done at P5 – P6 for developmental knockdown and at 4-7 weeks for adult knockdown. Injections were performed under a mixture of oxygen and isoflurane anesthesia (Baxter AG, Vienna, Austria) as previously described. The animals were placed in a stereotaxic frame (Kopf Instrument) and a single craniotomy was made over the VTA at following stereotaxic coordinates: ML +0.15 mm, AP +0.2 mm, DV −4.2 mm from lambda for P5-P6, and for 4-7 weeks: ML ±0.4 mm, AP −3.2 mm, DV −4.4 mm from Bregma. Injections were made with a 33-G Hamilton needle (Hamilton, 65460-02) for a total volume of 200 nL. Injections sites were confirmed *post-hoc* by immunostaining on VTA. The virus was incubated for 3 – 4 weeks prior to perform the behavioral tasks or immunostaining. Mice were excluded from the study if the body weight was less than 75% of the mean body weight at the start of behavior trials.

Injections of rAAV5-Ef1*α*-DIO-hChR2(H134R)-eYFP and rAAV5-Ef1*α*-DIO-eYFP were performed in DAT-Cre mice at 4 – 5 weeks. Mice were anesthetized and placed in a stereotactic frame (Angle One; Leica, Germany) as previously described. The skin was shaved, locally anesthetized with 40 – 50 µL lidocaine 0.5% and disinfected. Unilateral craniotomy (1 mm in diameter) was then performed to reach the VTA with a 10° angle, at following stereotactic coordinates: ML ±0.9 mm, AP –3.2 mm, DV – 4.20±0.05 mm from Bregma (Paxinos). The virus was injected via a glass micropipette (Drummond Scientific Company, Broomall, PA) into the VTA at the rate of 100 nl/min for a total volume of 500 nL. The virus was incubated for 3 – 4 weeks prior to perform the fiber optic implantation.

Implantations of optic fibers in VTA::DAChR^2^ and VTA::DAeYFP mice were then performed. Optic fibers were homemade built with Thorlabs materials, based on previously published protocol ^87^. Mice were anesthetized, placed in a stereotactic frame, the skin was shaved and a unilateral craniotomy was performed as previously described. The fiber optic was implanted with a 10° angle at the following coordinates: ML ±0.9 mm, AP –3.2 mm, DV –3.95±0.05 mm from Bregma above the VTA. The optic fiber was fixed on the skull using dental acrylic.

Implantations of stainless steel 26-gauge cannula (PlasticsOne, Virginia, USA) were performed on WT mice at 8 – 10 weeks. Mice were anesthetized and placed in a stereotactic frame as previously described. Unilateral craniotomy (1 mm in diameter) was then performed over the VTA at following stereotactic coordinates: ML ±0.9 mm, AP –3.2 mm, DV –3.95±0.05 mm from Bregma. The cannula was implanted with a 10° angle, placed above the VTA and fixed on the skull with dental acrylic. Between experiments, the cannula was protected by a removable cap. All animals underwent behavioral experiments 1 – 2 weeks after surgery.

### Immunohistochemistry and cell counting

VTA::DAhM^4^Di infected mice were deeply anesthetized and trans-cardially perfused with PBS 1× followed by 4% paraformaldehyde prepared in PBS 1×. The brain was removed and left for post-fixation at 4 °C in PBS 1×. Coronal VTA slices were cut at 50 µm and washed three times in PBS 1× before incubation with blocking solution containing 0.3% Triton X-100 and 1% goat serum. Slices were incubated with rabbit anti-TH (Abcam ab112, 1:500) at 4 °C overnight and then washed three times in PBS 1× and incubated for 2 hours at room temperature with secondary antibodies, goat anti-rabbit IgG-Alexa 488 (Abcam, 1:500; ab150077). Finally, the slices were washed three times in PBS 1× before being mounted onto microscope slides with Abcam DAPI mounting medium (Abcam, ab104139). Images were acquired with an LSM-700 confocal microscope.

Cell counting was performed on 50 µm thick VTA slices from 5 VTA::DAhM^4^Di. For each slice, images from the VTA and SNc were acquired bilaterally along the whole VTA dorso-ventral axis. The TH+, mCherry+ and TH+/mCherry+ cells were counted from different field of view. The total percentage of cells was calculated by averaging the total number of TH+ and TH+/mCherry+ of each mouse. The same procedure was performed for the SNc. An immunochemistry was performed for all the mice assess viral expression. Non-infected animals were excluded from the analysis.

VTA::DANL^3^KD mice were perfused as described above. Tissues were sectioned at 35 µm on a cryostat (Microm HM650, Thermo Scientific). Floating sections were kept in PBS 1× before incubation with blocking solution containing 0.5% Triton X-100 in TBS 1x and 10% normal donkey serum. The slices were incubated with sheep anti-TH (Millipor, AB1542, 1:1000) at 4 °C overnight and washed three times in 1x TBS containing 0.5% Triton X-100, followed by incubation for 2 hours at room temperature with a secondary antibody, donkey anti-sheep IgG-Cy3 (Jackson ImmunoResearch, 713-165-147, 1:1000). The sections were washed three times in TBS 1x containing 0.5% Triton X-100 before mounted onto microscope slides with ProLong Gold antifade (Invitrogen, p36930). Images were acquired on a custom-made dual spinning disk microscope (Life Imaging Services GmbH, Basel Switzerland) using 10x and 40x objectives. Images of brain regions expressing DAT; the VTA, SNc, dorsal raphe nucleus (DR) and retrorubral area (RR) were taken bilaterally along the whole dorso-ventral axis and images from at least 3 slices were counted. VTA::DANL^3^KD mice were included if a minimum of 20% of cells in the VTA were TH and GFP positive. Total percentage of infected cells was calculated by averaging the percentage obtained for each mouse.

### 3-chamber test

A three-chambered social preference test was used, comprising a rectangular Plexiglas arena (60 × 40 × 22 cm) (Ugo Basile, Varese, Italy) divided into three chambers (each 20 × 40 × 22 cm). These three chambers communicate by removable doors situated on the walls of the center chamber. 3 – 8 weeks after virus infusions, VTA::DAhM^4^Di and VTA::DAmCherry mice were randomly assigned to two batches and received intraperitoneal injection either with saline (vehicle) or Clozapine N-oxide (CNO, Enzo Life Science, Farmingdale, USA). All injections were done 30 minutes before starting the experiment. 1 – 2 weeks after, the mice that performed the task under CNO received vehicle and *vice versa*, therefore performing the task in both conditions. Each mouse was placed in the arena for 10 minutes of habituation during which it was free to explore the empty arena. At the end of the habituation, the mouse was temporarily kept in the center chamber by closing the removable doors. Two enclosures were placed in centers of the side chambers. One enclosure was left empty (inanimate object, o1) and the other one contained a novel social stimulus (novel juvenile mice C57BL/6J, 3 – 4 weeks, s1/s3 in vehicle or CNO condition). The doors were removed and the experimental mouse freely explored the arena and the two enclosures for 10 minutes. The walls of the enclosures, consisting of vertical metal bars, allowed visual, auditory, olfactory and tactile contact between the experimental mouse and the stimulus mouse. The stimuli mice were habituated to the enclosures during 3 sessions of 20 minutes the 3 days before the experiment. The position of the stimuli were randomly assigned and counterbalanced for every mouse.

The mice were then restrained a second time in the center chamber by closing the removable doors. The enclosures were held in position and a novel stimulus (s2/s4 in vehicle or CNO condition) was placed in the empty one. In this phase, the prior novel social stimulus is considered as familiar (social familiar, s1/s3). The doors were opened and the experimental mouse explored the arena for 10 minutes, with the two enclosures containing the familiar and the novel social stimuli. At the end of the 10 minutes, the experimental and stimuli mice returned to their home cage.

Every session was video-tracked and recorded using Ethovision XT (Noldus, Wageningen, the Netherlands), which provided the time in the different chambers and the distance moved during the test. An experimenter blind to the treatment of animals also manually scored behavior. The stimulus interaction was scored when the nose of the experimental was oriented toward the enclosures at a distance approximately less than 2 cm. The time interaction was used to calculate the Social Novelty Index as: *Interaction_Novel_ – Interaction_Familiar_*. The arena was cleaned with 5% ethanol solution and dried between trials.

### Intra-peritoneal injection of saline and Clozapine-N-oxyde (CNO)

The mice were weighted before each experiment and intra-peritoneal (i.p.) injection. The CNO dose was based on previous publications ^88^ and a concentration of 5mg/kg^−1^ was used for all the experiments. The CNO was, iluted in saline to obtain a concentration of 0.5mg.mL^−1^ to inject a reasonable volume of solution. The volume of saline (vehicle) injection was comparable to the volume of CNO solution.

### Long-term habituation/novelty exploration

An experimental cage similar to the animal’s home cage was used for this task. The bedding was cleaned after each trial and water and food were available.

During the habituation phase (4 days, day 1 to day 4), all VTA::DAhM^4^Di experimental mice (3 – 8 weeks after virus infection) received an intraperitoneal injection of saline 30 minutes before the task.

The experimental VTA::DAhM^4^Di mouse was placed in the cage with a novel social stimulus (juvenile mouse, C57Bl/6J, 3 – 4 weeks old, s1). The animals were let free to explore the cage and to interact with each other for 15 minutes. At the end of the trial, the experimental and stimulus mice were returned to their homecage. For 4 consecutive days the experimental mouse was exposed to the same social stimulus (s1) leading to a habituation of the environment and the social stimulus. Day 5 consisted in the novelty phase. The VTA::DAhM^4^Di experimental mice were split in two batches and were injected with either saline or CNO 30 minutes before the trial. A novel social stimulus (s2) was placed with the experimental mouse in the cage for 15 minutes to allow direct interaction. In total, the experimental mice were exposed to 2 different social stimuli: one social stimulus repeatedly presented from day 1 to day 4 (habituation phase, s1) and a second mouse at day 5 (novelty phase, s2) The same protocol as described above was used for object habituation/novelty. The VTA::DAhM^4^Di experimental mice received injection of saline and were exposed to the same object (der klein kaufman tanner; Germany, o1) from day 1 to day 4 (habituation phase). On day 5 the animals were injected with either saline or CNO, and were exposed to a novel object stimulus (novelty phase, o2).

To exclude pharmacological effects of the CNO dose on the behavioral parameters analyzed in this task, VTA::DAmCherry mice underwent the habituation/novelty and received an intraperitoneal injection of saline from day 1 to day 4 and CNO on day 5. The social and object habituation/novelty task performed with *Nlgn3KO* and VTA::DANL^3^KD was performed as described above. The test was done in a cage similar to the mice home cage containing food and water; the same cage was used for the duration of the trial. 3 – 4 weeks old C57Bl/6J male mice were used as stimulus mice, lego blocks and a small plastic toy were used as object. The animals were left to freely interact with the stimulus mouse or object for 15 minutes. For 4 consecutive days the experimental mouse was exposed to the same stimulus (s1 or o1). Day 5 consisted of the novelty phase (s2 or o2). At the end of each trial, the experimental and stimulus mice were returned to their home cage.

During the social habituation/novelty exploration task, non-aggressive social interaction was scored (experimenter blind to genotype and treatment group) when the experimental mouse initiated the action and when the nose of the animal was oriented toward the social stimulus mouse only. A non-aggressive interaction only initiated and maintained by the social stimulus mouse was not scored. During the object habituation/novelty exploration, the interaction was scored when the nose of the animal was oriented toward the object stimulus. The time interaction was used to calculate the Novelty Index as: *Interaction Day_5_–Interaction Day_4_*, both for social and object habituation/novelty exploration task.

The experimental cage was cleaned with 5% ethanol solution and the bedding was changed between sessions.

For the experiments with pharmacological agents, mice were cannulated to allow the infusion of either saline or 1–Naphthylacetyl spermine trihydrochloride (NASPM), directly in the VTA. The habituation task was performed as previously described. NASPM or saline were infused using a Minipump injector (pump Elite 11, Harvard apparatus, US) with 500 nL of saline (2 minutes of active injection at 250 nL/min rate, and 1 minute at rest), 10 minutes before each trial. At day 1, mice received saline. From day 2 to day 4 of the habituation phase, mice received either 4 µg of NASPM dissolved in 500 nL of saline or 500 nL of saline only (at 250 nL/min) before each trials. This dose has been previously used to obtain GluA2-lacking AMPARs block *in vivo*^89^. After at least 1 week, the animals were re-tested to habituation/novelty and the pharmacological treatment was counterbalanced. The scoring of the social or object interaction was made as previously described. The experimental cage was cleaned with 5% ethanol solution and the bedding was changed after every session. To assess the cannula placement, experimental subjects were infused using Chicago Sky Blue 6B (1 mg/mL), sacrificed 1 – 2 hours later and transcardially perfused as previously described.

For the experiments using optogenetic tools, VTA::DAChR^2^ and VTA::DAeYFP mice were implanted with a fiber optic above the VTA. All the mice underwent optogenetic bursting stimulation (5 pulses of 4ms at 20Hz with 500 ms between the beginning of each burst, ^34^) in a cage similar to home cage during 15 minutes. The optogenetic stimulation was non-contingent to the presence of a social stimulus. The mice used for electrophysiological recordings were sacrificed 24 hours after and the brain was sliced for *ex vivo* experiments, while the mice used for behavior started the habituation phase 24 hours after the 1st stimulation session. The mice underwent this stimulation protocol every day during 4 days (from day 0 to day 3), 5 hours after the free social exposure. Laser power was controlled between each test to ensure an estimated 7 – 10 mW of power at the implanted fiber tip. To assess the fiber placement and the viral infection, experimental subjects were sacrificed at the end of the habituation phase and transcardially perfused as previously described.

### Acute familiar exposure and Open Field with NASPM/saline

Mice were cannulated (at 8 – 10 weeks of age) and housed two per cage. After recovery (1 – 2 weeks), subjects were infused with either 500 nL of saline or 500 nL of NASPM (4µg/0.5µL at 250nL/min) 10 minutes before the trial. After infusion, mice returned to their home cage and the free non-aggressive interaction with their cage-mates was immediately scored for 15 consecutive minutes. 10 – 15 minutes after the social familiar exposure, the cannulated mice were placed in an open field arena for 10 minutes. The apparatus consisted in a 45 cm sided Plexiglas squared arena. After the test, experimental mice returned to their homecage. After 1 week the mice were re-tested and the animals that performed the task under NASPM received saline and *vice versa*. At the end, all the mice underwent both conditions.

The familiar social interaction was manually scored as described previously and the open field task was video-tracked (Ethovision, Noldus, Wageningen, the Netherlands) to automatically obtain the distance and the velocity during the session. The arena was cleaned with 5% ethanol solution after every session.

### Novelty conditioned place preference

Conditioned place preference experiments for examining reinforcing properties of novel social or object interactions were conducted in an apparatus (spatial place preference; BioSEB) consisting of two adjacent chambers (20 × 20 × 25 cm) with dots (black) or stripe (grey) wall patterns, connected by a lateral corridor (7 × 20 × 25 cm) with transparent walls and floor. The dots chamber was always associated to rough floor, while the stripe chamber with smooth floor. The illumination level was uniform between the two chambers and set at 10 – 13 lux. ANY-Maze behaviour tracking software was used to track animal’s movements within the apparatus and to manually score the time spent in non-aggressive interaction with the stimulus.

At day 0, experimental mice (male C57Bl6/J; group-housed; 8 – 16 weeks) freely explored the CPP apparatus for 15 minutes to determine Pre-TEST preference for one or the other chamber. After the Pre-TEST, experimental mice returned to their home cage with their cage-mates. The preference score was calculated as time spent in stimulus chamber (US+) divided by the sum of the time spent in stimulus chamber (US+) and the time spent in the empty chamber. No animals were excluded from the analysis based on preference score and US+ pairings were randomly assigned to dots or stripe chamber. At day 0, novel stimuli mice (male C57Bl/6J; single-housed; 3 – 4 weeks) or familiar stimuli mice (male C57Bl/6J; co-housed with experimental mice during the conditioning) were habituated to the US+ chamber for 15-30 minutes. The novel object stimulus was the same used in the object habituation/novelty task.

From day 1 to day 4, experimental mice underwent a conditioning schedule consisting of 30 minutes sessions (1 per day). Each session was subdivided in 6 blocks of 5 minutes during which the animals alternated between US+ and US-chamber, in presence (either familiar mouse, f1, novel mouse, s1 or novel object, o1) or absence (empty) of the stimulus, respectively. Experimental mice were guided through the corridor during the alternations and returned to their home cage with their cage-mates at the end of the conditioning session. Groups were counterbalanced for US+/US-sequences and for dots or stripes wall pattern. VTA::DAhM^4^Di and VTA::DAmCherry mice received an intraperitoneal injection of CNO (5 mg/Kg) 30-90 minutes *prior* each conditioning session.

At day 5, during the Post-TEST, experimental mice freely explored the CPP apparatus, without any stimulus for 15 minutes and the preference score was measured. The CPP apparatus was cleaned with 1% acetic acid, rinsed with distilled water and dried between each experimental subject.

### Social conditioned place preference

Mice were tested at P30 – P45 and were group housed before the test. The test apparatus was a custom-built cage measuring 46×24×22 cm divided into three chambers. The two outer chambers (23×18×22 cm) had vertical or horizontal striped pattern on the walls and flooring consisting or black rubber mats with different patterns (stripes *vs* squares). The outer chambers were joined together by a smaller chamber (23×10 cm) with white walls and floor with a 7×7 cm opening at the base to the outer chambers that can be closed. The cage was cleaned with 70% ethanol between each trial. During the pre-trial, mice were left to freely explore the cage for 30 minutes. After the pre-trial, all mice were single housed for the remainder of the test and one chamber was assigned the social chamber and one the isolation chamber. All mice received one social and one isolation condition session (30 minutes each) per day for 4 days, with a two-day rest between the 2nd and 3rd conditioning day. Mice were socially conditioned for 30 minutes together with their cage-mates followed by conditioning in the isolation chamber for 30 min. After the 4th conditioning day, mice were tested in a 30 minutes post-conditioning trial. The time spent freely exploring the chambers for 30 minutes was manually scored by an investigator blinded to the genotype. The preference score was calculated as the time spent in the social chamber divided by the combined time spent in the social and isolation chamber. Animals were excluded by pre-established criteria if they exhibited a strong preference for one chamber (more than 2x preference for one chamber).

### Olfactory habituation/dishabituation test

The olfactory habituation/dishabituation test was performed as previously described ^52^. Briefly, mice were individually tested for time spent sniffing cotton tipped swabs suspended from the cage lid. Distilled water, almond flavoring and banana flavoring (McCormick, Hunt Valley, MD; 1:100 dilution) and two different social odors were tested. Social odors were originated from two cages with the same number of male mice with different parental origins maintained for 6 days in the same bedding. Before the test, swabs were wiped in a zig-zag pattern across the bottom surface to collect the olfactory cues. Mice were acclimatized for 30 min with a cotton swab before testing. The order of presentation was: water, water, water, almond, almond, almond, banana, banana, banana, social odor 1, social odor 1, social odor 1, social odor 2, social odor 2, and social odor 2. Each swab was presented for a 2 min period, with a 1min interval between each presentation. Each test session was conducted in a clean mouse cage containing fresh litter. Time spent sniffing the swab was manuallyscored, the observers were blind of the genotype. Sniffing was scored when the nose was within 2 cm of the cotton swab. Mice were excluded by pre-established criteria if they did not investigate the first social odor (1 WT and 1 *Nlgn3KO* excluded).

### Open field, object recognition task, and marble burying

On day 1, mice were placed individually in the center of a square open field arena (50×50×30 cm) made of grey plastic for 7 minutes. Velocity (cm.sec^−1^) was analyzed using EthoVision10 system (Noldus). The arena was cleaned with 70% ethanol between trials. 24 hours later, mice were placed back in the arena containing two identical objects (culture flask filled with sand) for a 5-minutes acquisition trial. Object recognition memory was tested 1 hour later during a 5-minutes test trial in the arena containing a familiar and novel object (Lego block). The trial was recorded with a video camera and the time spent investigating was scored manually, the experimenters were blinded to the genotype. Investigation of the object was considered when the mouse nose was sniffing less than a centimeter from or touching the object. The discrimination ratio was calculated as following: 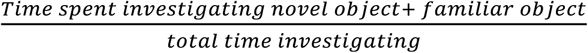. The arena and objects were cleaned with 70% ethanol between trials.

For the marble-burying test, animals were placed in a standard Type II cage with 5 cm bedding containing 20 identical black marbles distributed equally for 30 minutes. A marble was considered buried if at least 2/3 of the marble was covered.

### *Ex vivo* electrophysiology

200 – 250 µM thick horizontal midbrain slices were prepared from adolescence/early adulthood C57Bl/6J (4 – 16 weeks), VTA::DAhM^4^Di, VTA::DAmCherry, *Nlgn3* KO and VTA::DANL^3^KD mice. Subjects were anaesthetised with isoflurane/O2 and decapitated. Brains were sliced by using a cutting solution containing: 90.89 mM choline chloride, 24.98 mM glucose, 25 mM NaHCO3, 6.98 mM MgCl2, 11.85 mM ascorbic acid, 3.09 mM sodium pyruvate, 2.49 mM KCl, 1.25 mM NaH2PO4 and 0.50 mM CaCl2. Bain slices were incubated in cutting solution for 20-30 minutes at 35°. Subsequently, slices were transferred in artificial cerebrospinal fluid (aCSF) containing: 119 mM NaCl, 2.5 mM KCl, 1.3 mM MgCl2, 2.5 mM CaCl2, 1.0 mM NaH2PO4, 26.2 mM NaHCO3 and 11 mM glucose, bubbled with 95% O2 and 5% CO2) at room temperature. Whole-cell voltage clamp or current clamp electrophysiological recordings were conducted at 32°-34° in aCSF (2 – 3 ml.min^−1^, submerged slices). Recording pipette contained the following internal solution: 130 mM CsCl, 4 mM NaCl, 2 mM MgCl2, 1.1 mM EGTA, 5 mM HEPES, 2 mM Na2ATP, 5 mM sodium creatine phosphate, 0.6 mM Na3GTP, 0.1 mM spermine and 5 mM lidocaine N-ethyl bromide. *Ex vivo* CNO validation experiments were conducted in current-clamp configuration with the following internal solution: 140 mM K-Gluconate, 2 mM MgCl2, 5 mM KCl, 0.2 mM EGTA, 10 mM HEPES, 4 mM Na2ATP, 0.3 mM Na3GTP and 10 mM Creatine-Phosphate. Putative DA neurons of the VTA were identified accordingly to their position (medially to the medial terminal nucleus of the accessory optic tract), morphology, cell capacitance (> 28pF) and low input resistance at positive potentials. Excitatory post-synaptic currents (EPSCs) were recorded in voltage-clamp configuration, elicited by placing a bipolar electrode rostro-laterally to VTA at 0.1 Hz and isolated by application of the GABAAR antagonist picrotoxin (100 µM). The liquid junction potential was small (−3 mV) and traces were therefore not corrected. Access resistance (10 – 30 MΩ) was monitored by a hyperpolarizing step of −4 mV at each sweep, every 10 s. Data were excluded when the resistance changed > 20%. The AMPA/NMDA was calculated by subtracting to the mixed EPSC (+35 mV), the non-NMDA component isolated by D-APV (50 µM at +35 mV) bath application. The values of the ratio may be underestimated since it was calculated with spermine in the pipette. The rectification index (RI) of AMPARs is the ratio of the chord conductance calculated at negative potential (–60 mV) divided by the chord conductance at positive potential (+40 mV). The Paired Pulse Ratio (PPR) was measured at −60 mV, with a fixed inter stimulation interval of 50 msec. PPR was calculated by dividing the amplitude of the second EPSC by the amplitude of the first EPSC. To measure the RI from ChR2-expressing VTA DA neurons after *in vivo* optogenetic stimulation, a 500 msec blue-light pulse was delivered through the microscope objective in voltage-clamp configuration. Neurons were considered photocurrent positive (I +) when they responded with a large depolarizing current in response to optical stimulation, while photocurrent negative (I -) when they did not. Representative example traces are shown as the average of 10 – 20 consecutives EPSCs typically obtained at each potential. The synaptic responses were collected with a Multiclamp 700B-amplifier (Axon Instruments, Foster City, CA), filtered at 2.2 kHz, digitized at 5 Hz, and analyzed online using Igor Pro software (Wavemetrics, Lake Oswego, OR). Electrophysiology experiments were performed blind to behavioral or genetic manipulation.

### Drugs and viruses

rAAV5-hSyn-DIO-hM4D(Gi)-mCherry (Titer ≥ 3×10¹² vg.mL^−1^, Addgene), rAAV5-hSyn-DIO-mCherry (Titer ≥ 3×10¹² vg.mL^−1^, Addgene), rAAV5-Ef1*α*-DIO-hChR2(H134R)-eYFP (Titer ≥ 4.2×10¹² vg.mL^−1^, UNC Vector Core), rAAV5-Ef1*α*-DIO-eYFP (Titer ≥ 4.2×10¹² vg.mL^−1^, UNC Vector Core), pAAV2-Syn-DIO-miRNlgn3-GFP (Titer ≥ 7.2×109 vg.mL^−1^, miRNlgn3 RNAi BLOCK-iT RNAi Design, Invitrogen), pAAV2-Syn-DIO-miR-GFP (Titer ≥ 8.8×1011 vg.mL^−1^, RNAi BLOCK-iT RNAi Design, Invitrogen), pAAV2-Syn-GFP (Titer ≥ 1.4×1013 vg.mL^−1^), pAAV2-DIO-sh negative-GFP (Titer ≥ 8.8×1011 vg.mL^−1^, RNAi BLOCK-iT RNAi Design, Invitrogen), pAAV2-Syn-iCre (Titer ≥ 2.1×1012 vg.mL^−1^). Clozapine-N-oxyde (BML-NS105, Enzo), 1-Naphthylacetyl spermine trihydrochloride (24897539, Sigma-Aldrich), Picrotoxin (1128, Tocris) and D-APV (0106, Tocris).

### Cell culture

HEK293T (ATCC) transfected with V5-tagged *Nlgn3* and different NL3 knockdown plasmids (RNAi BLOCK-iT RNAi Design, Invitrogen) were maintained in DMEM supplemented with 10% FBS for 24 hours at 37°C after transfection. Cortical cultures were prepared from E16.5 mouse embryos. Neocortices were dissociated by addition of papain (130 units, Worthington Biochemical LK003176) for 30 min at 37°C. Cells were maintained in neurobasal medium (Gibco 21103-049) containing 2% B27 supplement (Gibco 17504-044), 1% Glutamax (Gibco 35050-038), and 1% penicillin/streptomycin (Sigma P4333). Neurons were transduced with recombinant AAV at DIV3 and maintained for 12-14 days. Viral transduction was performed in triplicates and viral knockdown assessed in ≥ 2 independent experiment.

### Biochemistry

Cortical neurons were lysed in lysis buffer containing 20mM Tris pH8.0, 100mM NaCl, 1mM EDTA, 1% Triton X-100, 0,2% SDS, 2mM DTT, and complete protease and phosphatase inhibitors (Roche Applied Science). Affinity-purified anti-NL3 and NL2 isoform-specific antibodies were previously described^47^. The following commercial available antibodies were used: mouse anti-tubulin (DHSB, Ab ID: AB_2315513 1:10000), mouse anti-synaptotagmin (Synaptic Systems, Cat. Nr. 105 011, 1:2000), rabbit anti-NeuN (Abcam, ab177487, 1:2000). Immunoblotting was done with HRP-conjugated secondary antibodies and Pierce ECL Western Blotting Substrate. Signals were acquired using an image analyzer (Bio-Rad, ChemiDoc MP Imaging System and Li-Cor, Odyssey) and images were analyzed using ImageJ.

### Statistical analysis

No statistical methods were used to predetermine the number of animals and cells, but suitable sample sizes were estimated based on previous experience and are similar to those generally employed in the field. The animals were randomly assigned to each group at the moment of viral infections or behavioral tests. Statistical analysis was conducted with GraphPad Prism 6 and 7 (San Diego, CA, USA) and MatLab (The Mathwork). Statistical outliers were identified with the ROUT method (Q=1) and excluded from the analysis. The normality of sample distributions was assessed with the Shapiro-Wilk criterion and when violated non-parametrical tests were used. When normally distributed, the data were analyzed with independent t-test, paired t-test, while for multiple comparisons one-way ANOVA and repeated measures (RM) ANOVA were used. When normality was violated, the data were analyzed with Mann-Whitney test, Wilcoxon matched-pairs signed rank test, while for multiple comparisons, Kruskal-Wallis or Friedman test were followed by Dunn’s test. For the analysis of variance with 2 factors (two-way ANOVA, RM two-way ANOVA and RM two-way ANOVA by both factors), normality of sample distribution was assumed, and followed by Bonferroni post-hoc test. All the statistical tests adopted were two-sided. When comparing two samples distributions similarity of variances was assumed, therefore no corrections were adopted. For social behavior experiments, the outlier analysis was conducted on manually scored non-aggressive social interaction. Data are represented as the mean ± s.e.m. and the significance was set at P < 0.05.

### Statistical Analysis of the 3-chamber task

According to the original developer of the 3-chamber task, this assay is a yes-or-no test in which animals display sociability/social novelty or they do not ^90^. For the statistical analysis and the interpretation of the results we followed the same principles, with minor adaptations. According to Nadler et al., 2004, during the sociability phase of the task, experimental subjects maintain higher time of interaction with a social stimulus compared to object for at least the first 10 minutes (as also evidenced in Bariselli, Tzanoulinou et al., 2016 by the maintenance of social preference across the 10 min-long social preference phase in control group). However, during the preference for social novelty, the experimental subjects maintain higher interaction time with a novel mouse compared to the familiar one only during the first 5 minutes of the test ^38^, while this difference is lost at later time points. For this reason, we decided to bin the time-course of both social preference and social novelty phases in 2 time intervals: first 5 minutes (0-5) and last five minutes (5-10). For the time in chamber analysis, we performed a RM one-way ANOVA on the time spent in social, center or object chamber or on time spent in novel, center of familiar stimulus chamber for the two time intervals, separately. If the ANOVA analysis gave a P < 0.05, we proceeded with multiple comparisons between of social *vs* center and social *vs* object (for the two time bins, separately) or novel *vs* center and novel *vs* familiar (for the two time bins, separately) by applying the Holm-Sidak correction. For the time sniffing, we performed a RM two-way ANOVA followed by Bonferroni post-hoc test. By performing this within-group analysis, we considered the animals to express sociability or preference for social novelty if they spent more time in the social/novel social chamber compared to the object/familiar mouse chamber and if they were engaged for longer time in social interaction or novel social interaction compared to object and familiar stimulus interaction, respectively^90^. Additionally, we calculated social novelty index, or preference score S2-S1 or S4-S3, to allow between group comparisons^42^ after a significant RM two-way ANOVA (P< 0.05 for main effects and interaction) followed by Bonferroni post-hoc test.

### Data availability

The data supporting this study are available upon request to the corresponding author.

### Statistical table

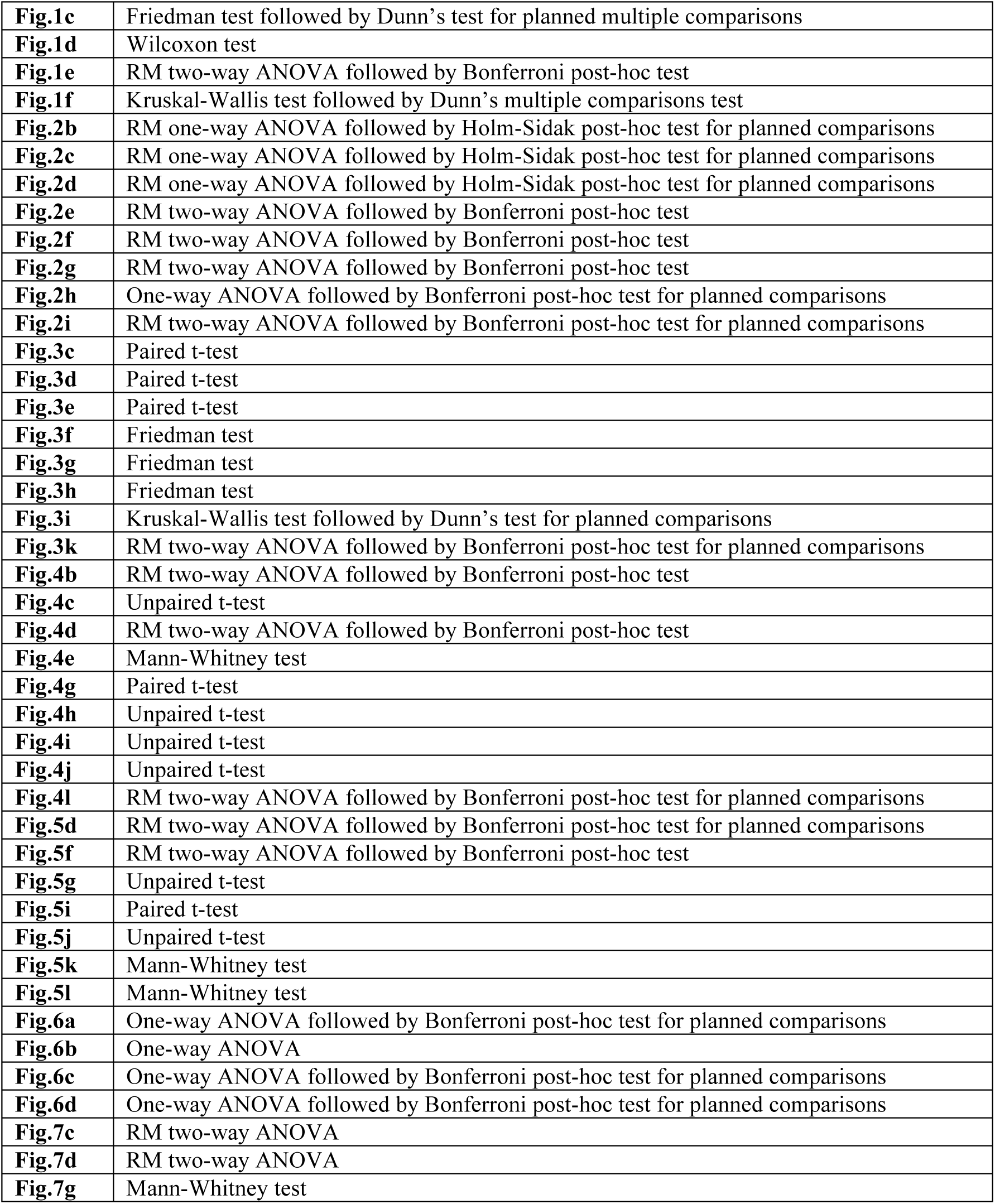
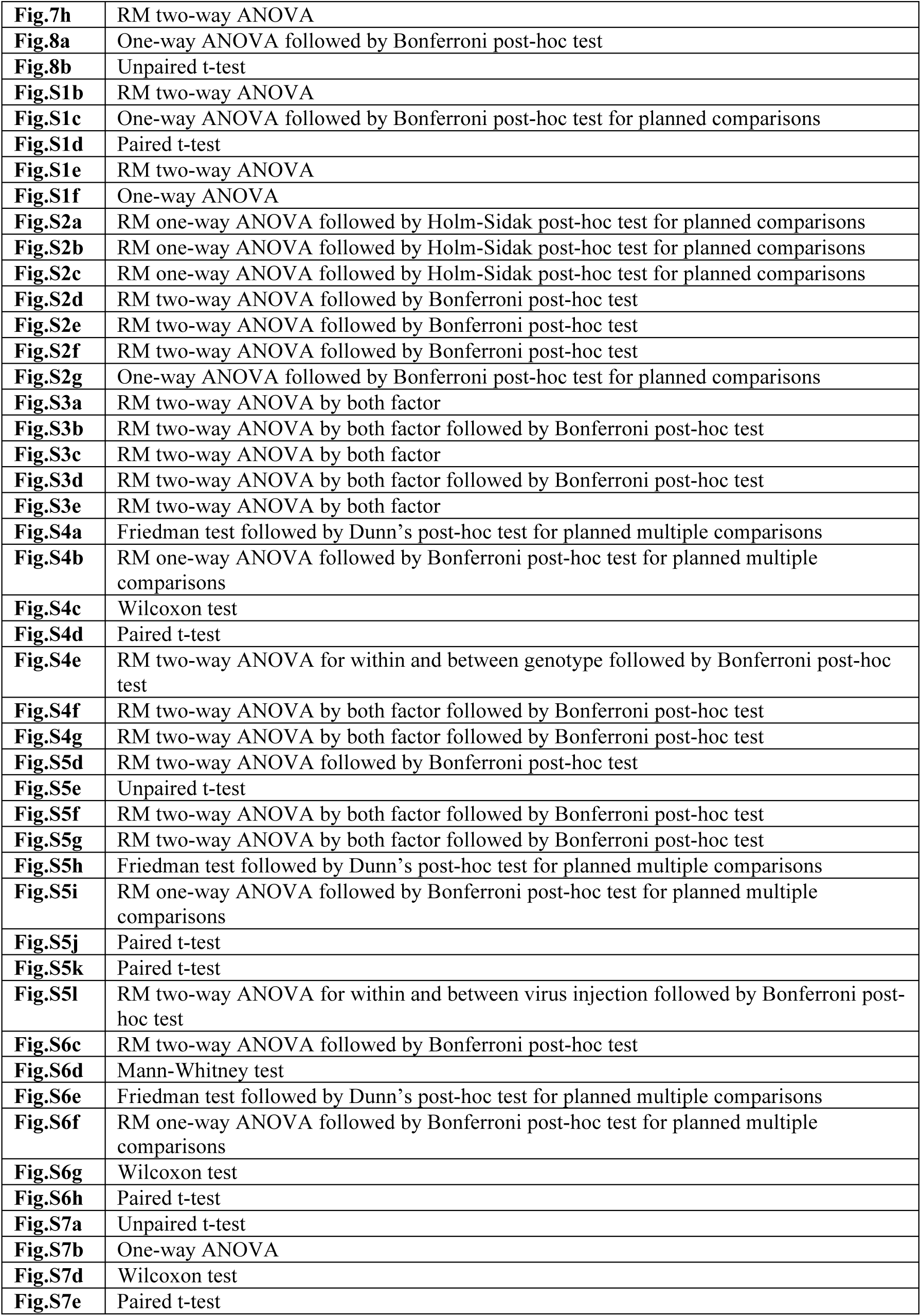

---

**Supplementary figure 1.**
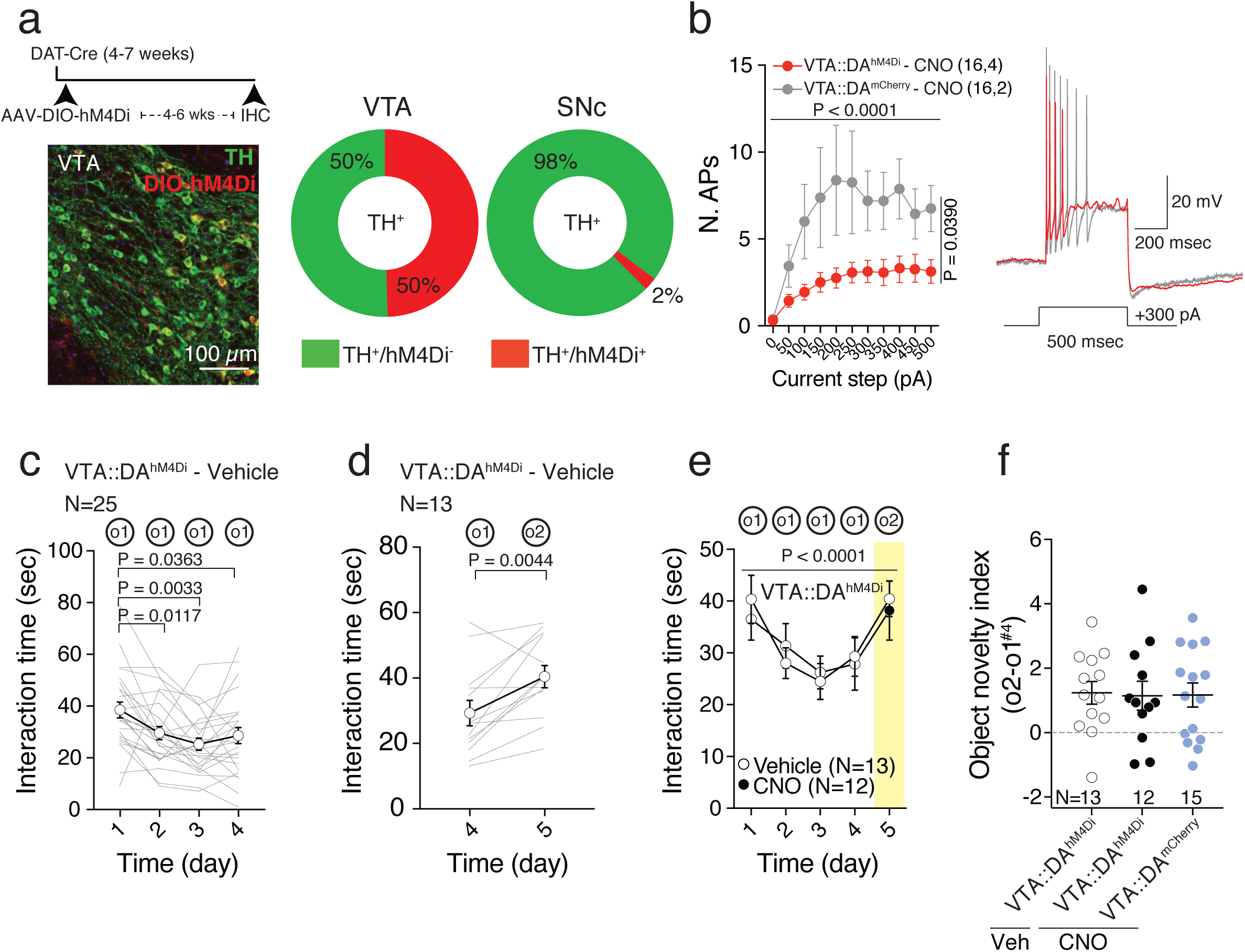
VTA DA neuron excitability controls social novelty exploration. **(a)** Top left: experimental paradigm. Bottom left: representative image of VTA DA neurons infected with AAV5-DIO-hM4Di-mCherry and stained against TH enzyme. Right: quantification of viral infection of TH+ neurons of VTA and Substantia Nigra pars compacta (SNc). **(b)** Left: graph representing the number of action potentials fired in response to 500 msec of increasing steps of current amplitude for VTA::DAhM^4^Di and VTA::DAmCherry neurons in presence of CNO (10 µM). RM two-way ANOVA (current main effect: F(10, 300) = 6.513, P < 0.0001; virus main effect: F(1, 30) = 4.66, P = 0.039; current × virus interaction: F(10, 300) = 1.527, P = 0.1287). Right: example traces of action potentials measured in current-clamp for VTA::DAhM^4^Di and VTA::DAmCherry neurons in CNO. **(c)** Time course of time interaction for VTA::DAhM^4^Di mice treated with vehicle. RM one-way ANOVA (F(2.31, 55.44) = 6.925, P = 0.0013) followed by Bonferroni post-hoc test for planned multiple comparisons. **(d)** Graph reporting the time interaction at day 4 with o1 and at day 5 with o2 for VTA::DAhM^4^Di mice treated with vehicle. Paired t-test (t(12) = 3.492). **(e)** Time interaction over days during habituation/novelty task for vehicle and CNO treated VTA::DAhM^4^Di mice (o1 and o2 are object novel stimuli presented at day 1-4 and day 5). RM two-way ANOVA (time main effect: F(4, 92) = 8.314, P < 0.0001; drug main effect: F(1, 23) = 0.01301, P = 0.9102; time × drug interaction: F(4, 92) = 0.4707, P = 0.7571). **(f)** Object novelty index calculated from VTA::DAhM^4^Di treated with vehicle, VTA::DAhM^4^Di and VTA::DAmCherry treated with CNO. One-way ANOVA (F(2,37) = 0.0144, P = 0.9857).

**Supplementary figure 2.**
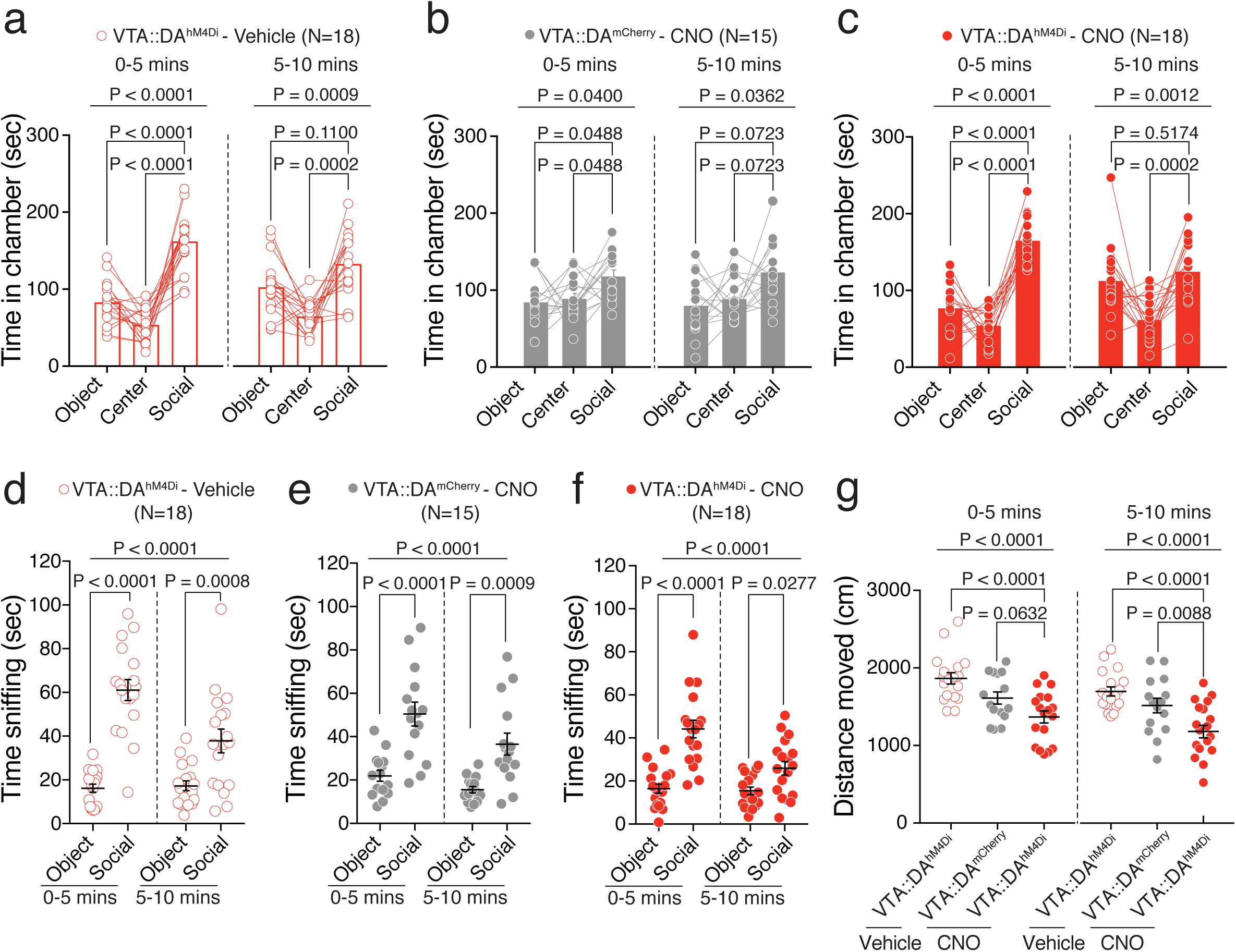
Effects of the reduction in VTA DA neuron excitability during social preference phase of the 3-chamber task. **(a)** Time in different stimuli chamber of vehicle treated VTA::DAhM^4^Di mice. RM one-way ANOVA binned in 5 mins (chamber main effect: F(1.519, 25.82) = 42.35, P < 0.0001, first 5 mins; chamber main effect: F(1.426, 24.25) = 11.63, P = 0.0009, last 5 mins) followed by Holm-Sidak post-hoc test. **(b)** Time in different stimuli chamber of CNO treated VTA::DAmCherry mice. RM one-way ANOVA binned in 5 mins (chamber main effect: F(1.857, 26) = 3.748, P = 0.0400, first 5 mins; chamber main effect: F(1.725, 24.15) = 4.026, P = 0.0362, last 5 mins) followed by Holm-Sidak post-hoc test. **(c)** Time in different stimuli chamber of CNO treated VTA::DAhM^4^Di mice. RM one-way ANOVA binned in 5 mins (chamber main effect: F(1.777, ^30^.^21^) = 56.17, P < 0.0001, first 5 mins; chamber main effect: F(1.653, 28.11) = 9.611, P = 0.0012, last 5 mins) followed by Holm-Sidak post-hoc test. **(d)** Time sniffing toward social or object stimuli of vehicle treated VTA::DAhM^4^Di mice. RM two-way ANOVA (stimulus main effect: F(1, 34) = 43.28, P < 0.0001; time main effect: F(1, 34) = 21.54, P < 0.0001; time × stimulus interaction: F(1, 34) = 25.97, P < 0.0001) followed by Bonferroni post-hoc test. **(e)** Time sniffing toward social or object stimuli of CNO treated VTA::DAmCherry mice. RM two-way ANOVA (stimulus main effect: F(1,28) = 23.65, P < 0.0001; time main effect: F(1,28) = 17.22, P = 0.0003; time × stimulus interaction: F(1,28) = 2.311, P = 0.1397) followed by Bonferroni post-hoc test. **(f)** Time sniffing toward social or object stimuli of CNO treated VTA::DAhM^4^Di mice. RM two-way ANOVA (stimulus main effect: F(1,34) = 34.13, P < 0.0001; time main effect: F(1,34) = 14.14, P = 0.0006; time × stimulus interaction: F(1,34) = 11.22, P = 0.0020) followed by Bonferroni post-hoc test. **(g)** Distance moved during the social preference phase, binned in 5 mins, for VTA::DAhM^4^Di vehicle and CNO treated mice, and VTA::DAmCherry CNO treated mice. One-way ANOVA (group main effect: F(2, 48) = 11.26, P < 0.0001, first 5 mins; group main effect: F(2, 48) = 12.03, P < 0.0001, last 5 mins) followed by Bonferroni post-hoc test for planned comparisons. N indicates number of mice. Error bars report s.e.m.

**Supplementary figure 3.**
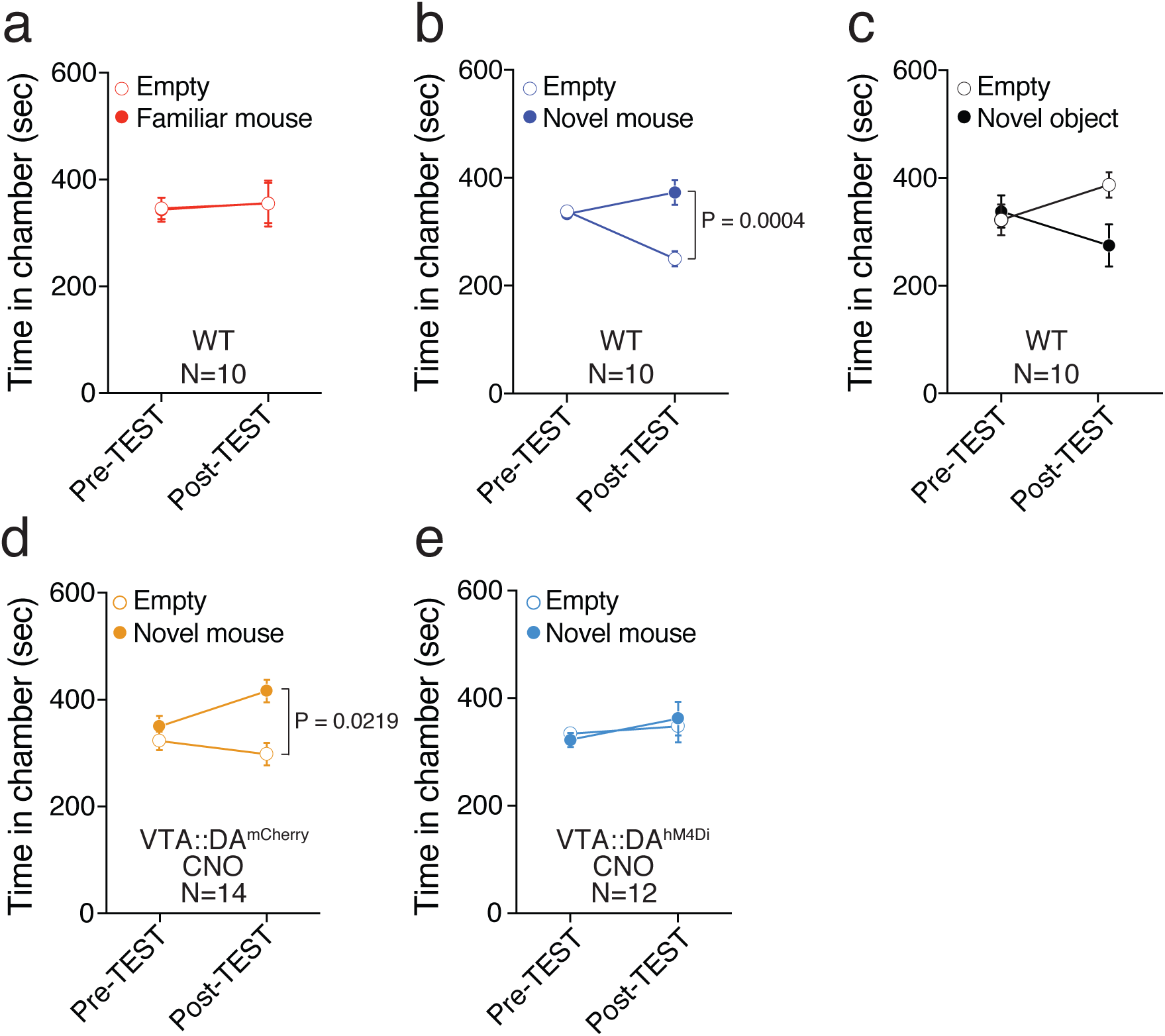
Time of chamber exploration during nCPP. **(a)** Graph representing the time spent in either empty or familiar mouse paired chamber during the Pre-and the Post-TEST. RM two-way ANOVA by both factors (time main effect: F(1, 9) = 1.499, P = 0.2519; chamber main effect: F(1, 9) = 0.0002, P = 0.9885; time × chamber interaction: F(1, 9) = 0.0033, P = 0.9557). **(b)** Graph representing the time spent in either empty or novel mouse paired chamber during the Pre-and the Post-TEST. RM two-way ANOVA by both factors (time main effect: F(1, 9) = 11.45, P = 0.0081; chamber main effect: F(1, 9) = 5.313, P = 0.0466; time × chamber interaction: F(1, 9) = 19.4, P = 0.0017) followed by Bonferroni post-hoc test. **(c)** Graph representing the time spent in either empty or novel object paired chamber during the Pre-and the Post-TEST. RM two-way by both factors (time main effect: F(1, 9) = 0.0098, P = 0.9232; chamber main effect: F(1, 9) = 0.9655, P = 0.3515; time × chamber interaction: F(1, 9) = 3.912, P = 0.0793). **(d)** Graph representing the time spent in either empty or novel social stimulus paired chamber during the Pre-and the Post-TEST for VTA::DAmCherry animals treated with CNO. RM two-way ANOVA by both factors (time main effect: F(1, 13) = 0.7315, P = 0.4002; chamber main effect: F(1, 13) = 26.9361, P < 0.0001; time × chamber interaction: F(1, 13) = 3.5527, P = 0.0707) followed by Bonferroni post-hoc test. **(e)** Graph representing the time spent in either empty or novel social stimulus paired chamber during the Pre-and the Post-TEST for VTA::DAhM^4^Di animals treated with CNO. RM two-way ANOVA by both factors (time main effect: F(1, 11) = 1.7620, P = 0.1980; chamber main effect: F(1, 11) = 0.0040, P = 0.9503; time × chamber interaction: F(1, 11) = 0.4080, P = 0.5296). N indicates number of mice. Error bars report s.e.m.

**Supplementary Figure 4.**
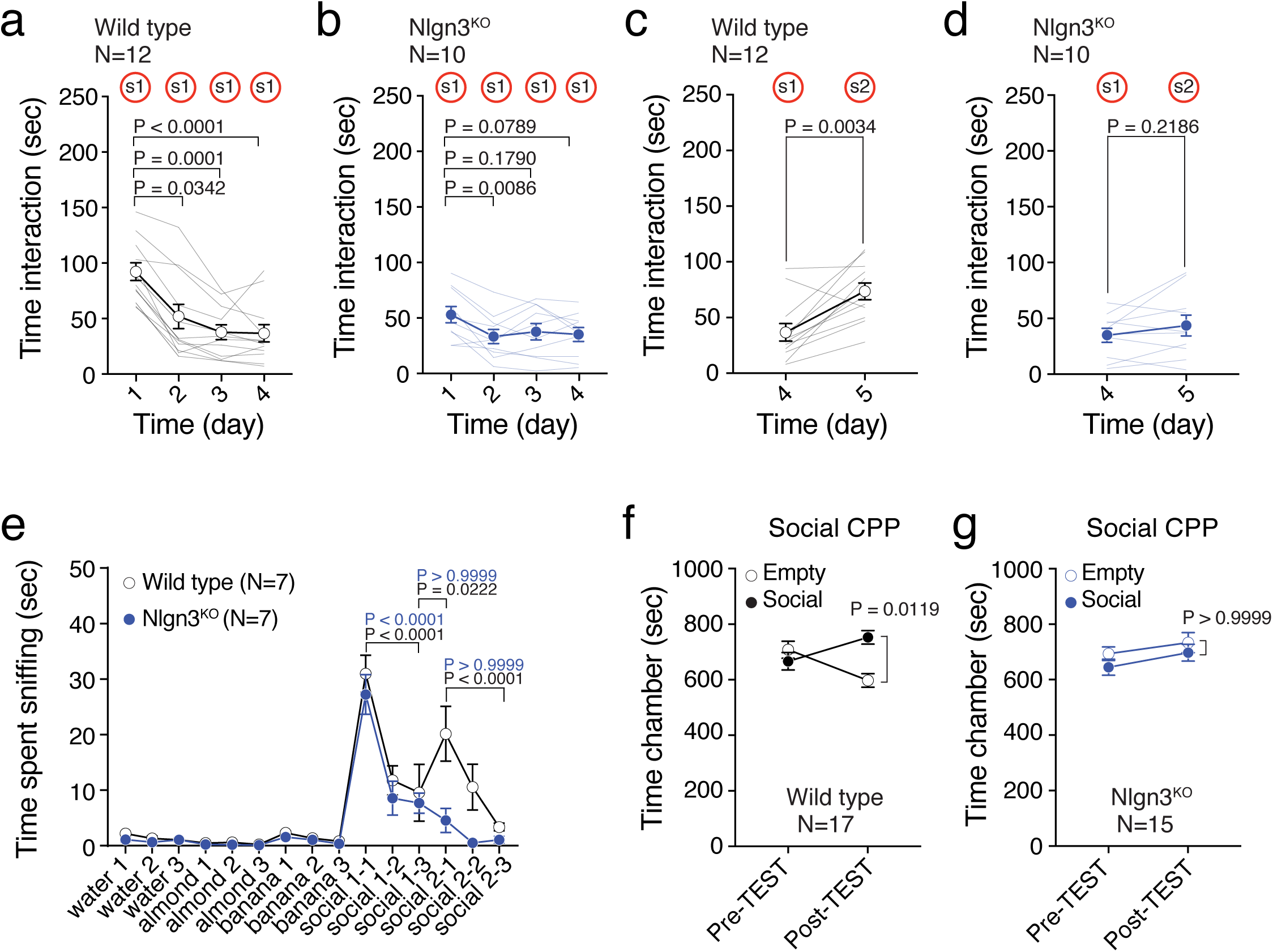
Global knockdown of *Nlgn3* alters social novelty and social reward response. (**a-b)** Time spent interacting with the stimulus mouse during day 1-4 of the social novelty exploration test plotted for (**a**) WT (Friedman test (x^2^(4) = 26.8, P < 0.0001) followed by Dunn’s post-hoc test for planned multiple comparisons) and (**b**) *Nlgn3KO* mice (RM ANOVA (F(2.054, 18.48) = 5.179, P = 0.0158) followed by Bonferroni post-hoc test for planned multiple comparisons). (**c-d**) Time interacting with s1 on day 4 and s2 on day 5 plotted for (**c**) WT (Wilcoxon test (W = 70) and (**d**) *Nlgn3KO* mice (Paired t-test (t(9) = 1.323). (**e**) Mean time spent sniffing the odor plotted for WT and *Nlgn3KO* mice. RM two-way ANOVA within genotype to evaluate habituation (P-values displayed in graph) and between genotype to evaluate differences in response (social 2-1: WT vs *Nlgn3KO* P<0.0001, social 2-2: WT vs *Nlgn3KO* P=0.0064) (odor main effect: F(14, 168) = 30.42, P < 0.0001; genotype main effect: F(1, ^12^) =15.42, P = 0.0020; odor × genotype interaction: F(14, 168) = 2.45, P = 0.0036) followed by Bonferroni’s post-hoc test. (**f-g**) Time spent in empty or social chamber during the social CPP Pre-and Post-TEST plotted for (**f**) WT (Two-way ANOVA with repeated measures for both factors (time main effect: F(1,16) = 1.361, P = 0.2604; chamber main effect: F(1,16) = 2.22, P = 0.1557; time × chamber interaction: F(1,16) = 8.07, P = 0.0118) followed by Bonferroni post-hoc test) and (**g**) *Nlgn3KO* (Two-way ANOVA with repeated measures for both factors time main effect: F(1,14) = 6.992, P = 0.0192; chamber main effect: F(1,14) = 1.233, P = 0.2855; time × chamber interaction: F(1,14) = 0.0225, P = 0.8827) followed by Bonferroni post-hoc test). N indicates number of mice. Error bars report s.e.m.

**Supplementary Figure 5.**
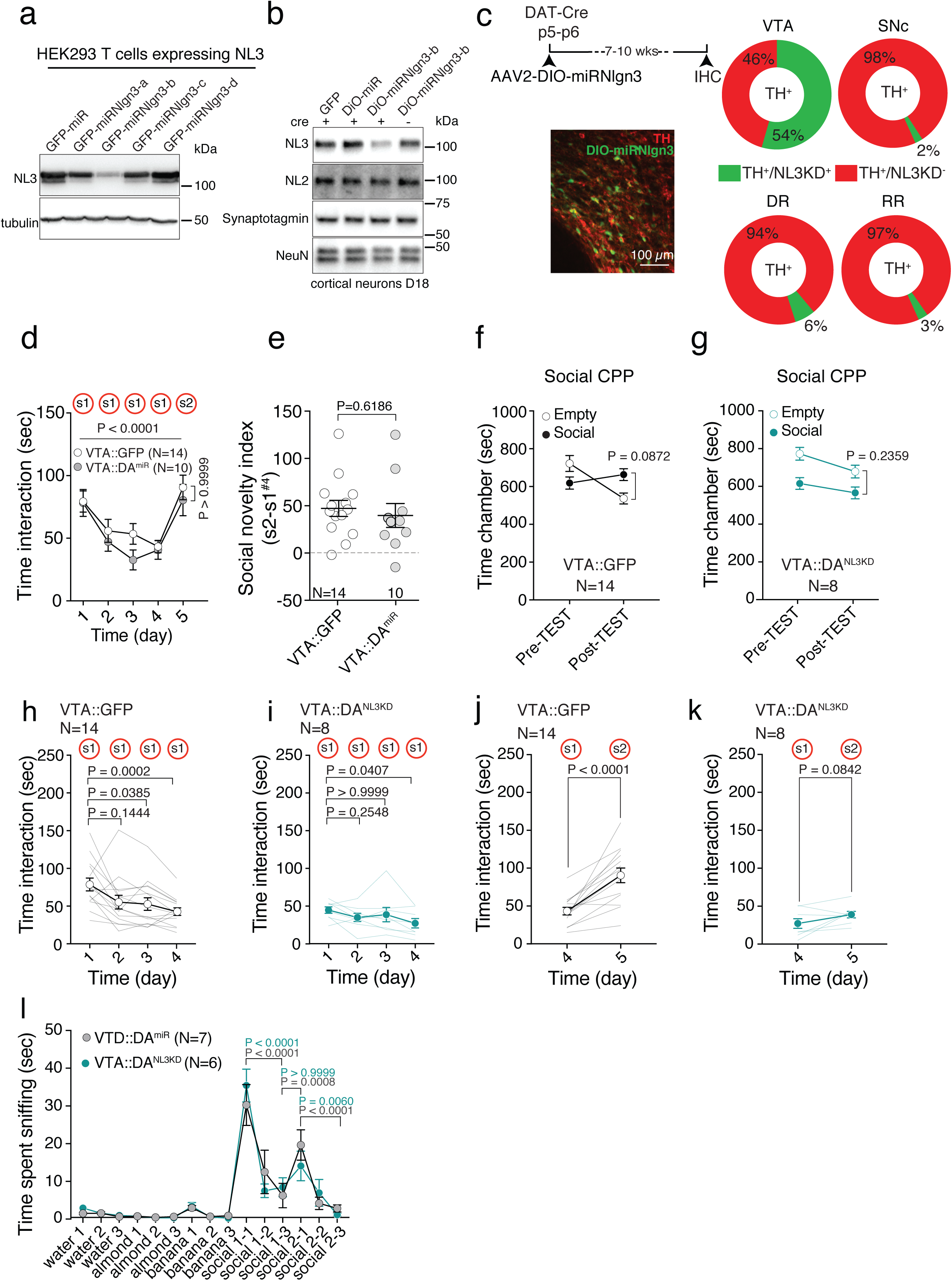
*Nlgn3* in VTA DA neurons is required for social novelty and social interaction. (**a**) Western blot of HEK293T cells expressing V5-tagged *Nlgn3* transfected with different *Nlgn3* miR knockdown construct. (**b**) Neuroligin 3 (NL3), Neuroligin 2 (NL2), synaptotagmin and NeuN protein levels in cortical neurons transfected with AAV2-Synaptophysin-GFP, AAV2-DIO-miR*Nlgn*^*3*^ or AAV2-DIO-miR. Protein abundance was probed by western blot from cortical neuron lysate collected at DIV18. (**c**) Top: experimental timeline for in vivo knock-down of NL3 expression in VTA-DA neurons. Left: Representative image of confocal section stained with anti-TH antibodies and visualizing GFP fluorescence driven from the AAV2-DIO-miRNlgn^3^. Right: Quantification of viral infection in TH-positive neurons in the VTA, SNc, dorsal raphe nucleus (DR) and retrorubral area (RR) of DA*TiCre* mice. TH+/ NL3KD+: double-positive neurons; TH+/NL3KD-: TH-positive GFP-negative neurons. Percentage represent mean from 8 animals. (**d**) A virus expressing a non-targeting, cre-dependent miR injected into DA*TiCre* mice (VTA::DAmiR) was compared to the VTA::GFP injected mice in the social novelty exploration task. Graph is reporting the mean social interaction. RM two-way ANOVA (time main effect: F(4, 88) = 19.42, P < 0.0001, virus main effect: F(1, 22) = 0.8246, P = 0.3737; time × virus interaction: F(4, 88) = 0.6914, P = 0.5998) followed by Bonferroni’s post-hoc test. (**e**) Social novelty index for VTA::GFP and VTA::DAmiR mice. Unpaired t-test (t(22) = 0.505)). This demonstrates that a non-targeting miR does not produce a phenotype in this assay. (**f-g**) Time spent in empty or social chamber during the social CPP Pre-and Post-TEST plotted for (**f**) VTA::GFP (Two-way ANOVA with repeated measures for both factors. Time main effect: F(1,13) = 9.285, P = 0.0094; chamber main effect: F(1,13) = 0.06717, P = 0.7996; time × chamber interaction: F(1,13) = 8.26, P = 0.0130) followed by Bonferroni post-hoc test) and (**g**) VTA::DANL^3^KD mice. Two-way ANOVA with repeated measures for both factors. Time main effect: F(1,7) = 8.405, P = 0.0230; chamber main effect: F(1, 7) = 14.49, P = 0.0067; time × chamber interaction: F(1,7) = 0.2515, P = 0.6314) followed by Bonferroni post-hoc test. (**h-i**) Time spent interacting with the stimulus mouse during day 1-4 of the social novelty exploration test plotted for (**h**) VTA::GFP (Friedman test (x^2^(4) = 16.8, P = 0.0008) followed by Dunn’s post-hoc test for planned multiple comparisons) (**i**) and VTA::DANL^3^KD mice. RM ANOVA (F(1.48, 10.36) = 3.098, P = 0.0978) followed by Bonferroni post-hoc test for planned multiple comparisons. (**j-k**) Time interacting with s1 on day 4 and s2 on day 5 plotted for (**j**) VTA::GFP (Paired t-test (t(13) = 5.537) and (**k**) VTA::DANL^3^KD mice (Paired t-test (t(7) = 2.011). (**l**) Mean time spent sniffing the odor plotted for VTA::DAmiR and VTA::DANL^3^KD mice. RM two-way ANOVA within virus injection to evaluate habituation (P-values displayed in graph) and between virus injection to evaluate differences in response (odor main effect: F(14, 154) = 34.26, P < 0.0001; virus main effect: F(1, 11) =0.0026, P = 0.9602; odor × genotype interaction: F(14, 154) = 0.7935, P = 0.6752) followed by Bonferroni’s post-hoc test. N numbers indicate mice. Error bars show s.e.m.

**Supplementary Figure 6.**
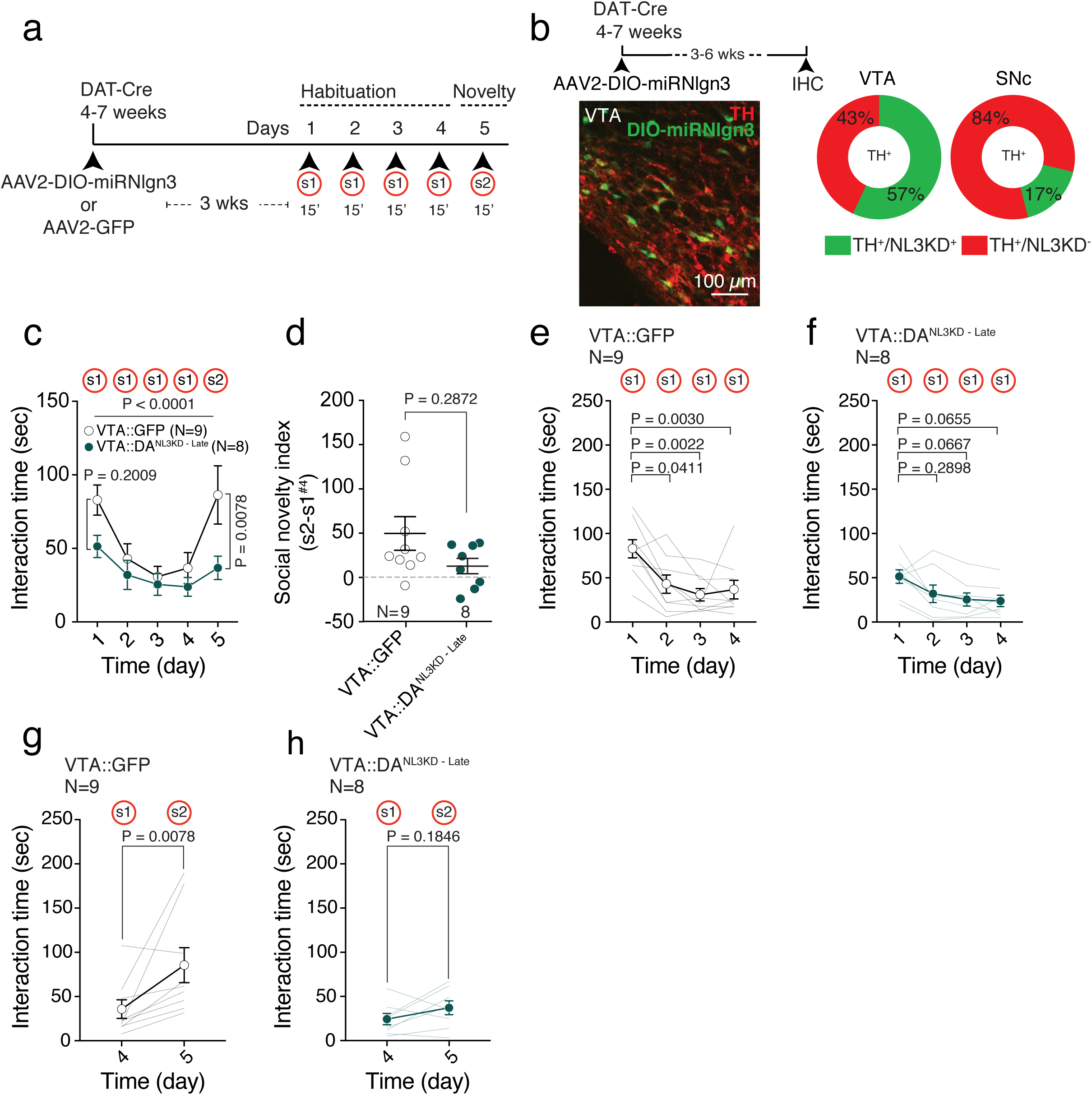
Adult *Nlgn3* knockdown in VTA DA neurons alters social novelty response. (**a**) Experimental schematics in adult VTA-injected mice. (**b**) Top: experimental timeline for in vivo knock-down of NL3 expression in adult VTA-DA neurons. Left: Representative image of confocal section stained with anti-TH antibodies and visualizing GFP fluorescence driven from the AAV2-DIO-miRNlgn^3^. Right: Quantification of viral infection in TH-positive neurons in the VTA and SNc of DA*TiCre* mice. TH+/ NL3KD+: double-positive neurons; TH+/NL3KD-: TH-positive GFP-negative neurons. Percentage represent mean from 8 animals. (**c**) Mean social interaction plotted for VTA::GFP and VTA::DANL^3^KD adult-injected mice. RM two-way ANOVA (time main effect: F(4, 60) = 10.18, P < 0.0001, virus main effect: F(1, 15) = 3.91, P = 0.0667; time × virus interaction: F(4, 60) = 2.579, P = 0.0463) followed by Bonferroni’s post-hoc test. (**d**) Social novelty index for adult-injected VTA::GFP and VTA::DANL^3^KD mice. Mann-Whitney U=24.5. (**e-f**) Time spent interacting with the stimulus mouse during day 1-4 of the social novelty exploration test plotted for (**e**) adult-injected VTA::GFP (Friedman test (x^2^_(4)_ = 15.07, P = 0.0018) followed by Dunn’s post-hoc test for planned multiple comparisons) (**f**) and adult-injected VTA::DANL^3^KD mice. RM ANOVA (F(1.6, 11.2) = 5.837, P = 0.0228) followed by Bonferroni post-hoc test for planned multiple comparisons. (**g-h**) Time interacting with s1 on day 4 and s2 on day 5 plotted for (**g**) adult-injected VTA::GFP (Wilcoxon (W= 43) and (**h**) adult-injected VTA::DANL^3^KD mice (Paired t-test (t(7) = 1.472). N numbers indicate mice. Error bars show s.e.m.

**Supplementary Figure 7.**
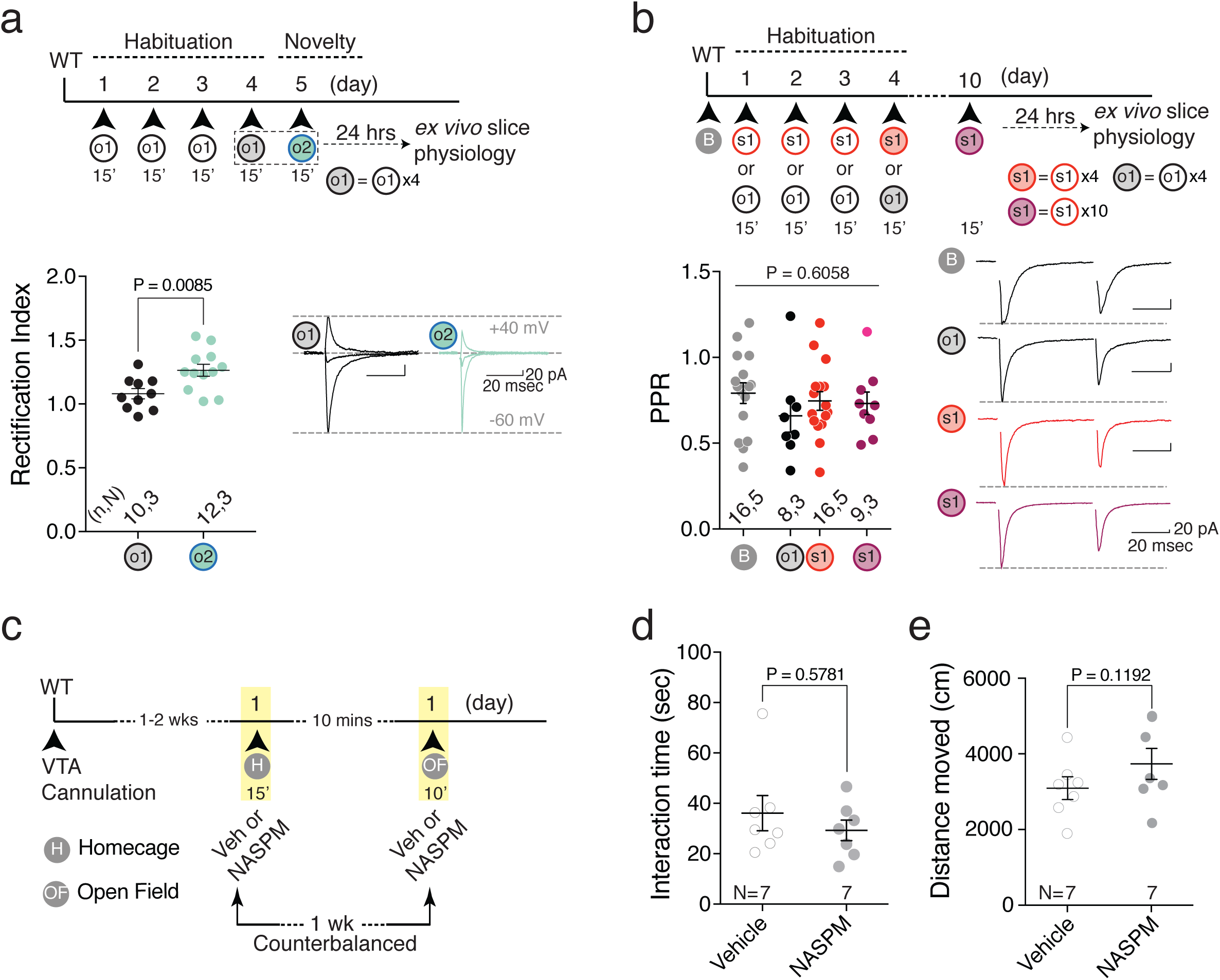
Additional characterization of synaptic and behavioural parameters during long-term habituation task. **(a)** Top: experimental paradigm. Bottom: scatter plot and example traces of rectification index measured from VTA DA neurons 24 hours after 4 repeated exposures to object novelty (o1) or 24 hours after exposure to novel object (o2) after 4 days exposure to o1. The data reported for o1 are the same used in Figure 6c. Unpaired t-test (t(20) = 2.918). (**b**) Top: experimental paradigm. Bottom: scatter plot and example traces of Paired-Pulse Ratio (PPR, −60 mV) measured 24 hours after 4 repeated exposures to either novel object (o1) or novel mouse (s1, red) and 24 hours after 10 repeated exposures to novel mouse (s1, purple). One-way ANOVA (F(3, 45) = 0.6058, P = 0.6147). **(c)** Experimental paradigm of cage mate interaction in home cage and distance travelled in open field of vehicle and NASPM VTA-infused mice. Mice received both vehicle and NASPM in each condition with 1 week interval. **(d)** Scatter plot of time interaction measured for 15 minutes in the home cage with cage mate of mice infused with either vehicle or NASPM. Wilcoxon (W= −8). **(e)** Scatter plot of distance travelled in open-field for 10 minutes of mice infused with either vehicle or NASPM. Paired t-test (t(6) = 1.816).

